# A *De Novo*, Intragenic Modifier in *Cis* Rescues *Scn8a*-N1768D Epilepsy and Defines a Therapeutic Window Bounded by Gain- and Loss-of-Function

**DOI:** 10.64898/2026.09.08.750161

**Authors:** Erfan Bahramnejad, Victor J. Hiller, Michael F. Hammer

**Affiliations:** BIO5 Institute, University of Arizona, Tucson, AZ, U.S.A.; Department of Neurology, University of Arizona, Tucson, AZ, U.S.A

**Keywords:** frameshift, pathogenic variant, pediatric epilepsy, genetic modifier, goldilocks gene, therapeutic window, RNA-seq, gene expression, whole-genome sequencing, sodium channel

## Abstract

Gain-of-function variants in *SCN8A*, encoding Na_V_1.6, cause developmental and epileptic encephalopathy with early-onset, drug-resistant seizures. *SCN8A* is a “Goldilocks” gene: both excess and deficit of Na_V_1.6 are deleterious, so therapies that lower Na_V_1.6 must operate within a narrow window. Here we characterize a congenic C3H/HeJ.C57BL/6J colony carrying *Scn8a*-N1768D and identify a tightly linked, *cis*-acting modifier (L) that converts a severe phenotype into graded outcomes. Among N1768D homozygotes on a 98.4% C3H/HeJ.C57BL/6J background, three phenotypes segregate: short-lived (SL) mice with early lethal tonic-clonic seizures (TCS), long-lived (LL) mice with delayed onset and reduced seizure burden, and hindlimb-paralysis (HP) mice with early death. Heterozygotes separate into D/+ SL mice, which develop spontaneous TCS and die of epilepsy, and D/+ LL mice, which lack spontaneous TCS and epilepsy-related death. Genetic analysis places a second locus (L) tightly linked to *Scn8a* and co-segregating with phenotype. Hippocampal RNA-seq shows dose-dependent *Scn8a* mRNA reduction: none in SL (D+/D+), ∼30% in LL (DL/D+), and ∼70% in HP (DL/DL). Long-read trio DNA sequencing identifies a *de novo* 2-bp frameshift in coding exon 15 of *Scn8a*, in *cis* with N1768D, that introduces a premature stop codon ∼26 kb upstream of the 3′-most exon–exon junction, predicted to trigger nonsense-mediated decay. Allele-specific read counting confirms selective, *cis*-acting loss of DL transcripts with preserved D+ output. Thus, a *de novo*, intragenic frameshift variant titrates Na_V_1.6 through a Goldilocks window in which intermediate reduction rescues Gain-of-function epilepsy, whereas insufficient or excessive reduction remains lethal, validating *Scn8a*-lowering and its therapeutic dose window.

## 1. Introduction

Brain-expressed voltage-gated sodium channels (Na_V_) are essential for maintaining the balance of neuronal excitability, playing a key role in membrane potential and neuronal firing. Genetic alterations in these channels can disrupt this balance, leading to epilepsy and/or developmental impairments through gain-of-function (GoF) mechanisms (1). GoF variants in Na_V_ channels enhance neuronal firing through premature channel opening, increased persistent current, or delayed inactivation, leading to hyperexcitability and seizures. For example, pathogenic variants with GoF effects at three of the four brain-expressed channels (*SCN2A*, *SCN3A* and *SCN8A*) are all associated with pediatric epilepsy (2). Patients with GoF variants in *SCN8A* present with a spectrum of phenotypes, including mild-to-severe developmental and epileptic encephalopathy (DEE) and pharmacoresistant epilepsy with multiple seizure types, whereas those with pathogenic loss-of-function (LoF) variants present with neurodevelopmental delay with or without generalized epilepsy (3). Many of the features of DEE are mimicked in transgenic mouse models (2, 4).

Transgenic mouse models provide essential tools for dissecting these mechanisms and developing targeted therapies for genetic epilepsies, for which there is an urgent need to better understand disease mechanisms and identify more effective treatments. Unfortunately, up to 78% of patients with *SCN8A*-related epilepsy (*SCN8A*-RE) remain drug-resistant, a percentage that is almost three times higher than typical for other forms of pediatric epilepsy (5, 6). Genetic modifiers are defined as genes that alter the phenotypic expression of a primary disease-causing gene (7), with the potential to either exacerbate or ameliorate the effects of the primary mutation. Typically, modifiers do not cause disease on their own but interact with the disease gene or its products (8), influencing pathways that can affect, for example, ion channel function, neuronal excitability, or cellular responses to stress (9, 10).

An active area of pre-clinical research focuses on developing genetic therapies that reduce Na_V_1.6 expression to a level that mitigates hyperexcitability and seizure generation. One of the main challenges arises because *SCN8A* acts as a “Goldilocks” gene: both GoF and LoF variants disrupt normal physiology. This dual sensitivity creates a major barrier to therapy as treatments must preserve Na_V_1.6 activity within a narrow “functional window” (6). Defining this optimal level of Na_V_1.6 is critical. We have developed a series of transgenic mouse lines that carry the *Scn8a*-N1768D (D/+) GoF variant on the C57BL/6J (B6) and C3H/HeJ (C3H) backgrounds, representing a model system for studying natural genetic modifiers of SCN8A-RE. These lines emerged from a breeding program initiated to investigate the protective effects of C3H genetic background on B6 D/+ mice. We discovered a genetic locus that maps within 1.88 cM of the *Scn8a* gene on chromosome 15 and reduces *Scn8a* expression by approximately 30% (in long-lived heterozygotes) to 70% (in hindlimb-paralysis homozygotes) relative to wild-type levels in a dose-dependent manner and rescues the severe epilepsy phenotype of homozygous D/D mice. Here we present evidence that the modifier acts in an allele-specific manner with zygosity at this locus corresponding to three distinct phenotypes of D/D mice: short-lived mice with no modifier copies (SL; D+/D+), long-lived mice with one copy (LL; DL/D+), and hindlimb-paralysis mice with two copies (HP; DL/DL), the last exhibiting early death. Determining the mechanism by which this “natural genetic modifier” regulates *Scn8a* expression levels has the potential to inform therapeutic windows for emerging gene therapies and to define critical parameters of the “Goldilocks effect”.

## 2. Materials and Methods

### 2.1 Animals

The *Scn8a*-N1768D knock-in mutation was created using TALEN technology. The procedure was performed on fertilized eggs from a cross between B6 and SJL mouse strains (F2 generation) at the University of Michigan’s Transgenic Animal Model Core facility, following established protocols, which is fully explained in (11). We characterized the lifespan, age of onset, and seizure frequency of heterozygous (D/+) and homozygous (D/D) mice in B6 background and published our results. D/D mice in B6 background live around 24 days (12).

To characterize the same parameters of D/D mice on the C3H background, we created a congenic C3H.B6-*Scn8a*^N1768D^ (C3H.B6) line by backcrossing a male heterozygote (D/+) in B6 colony to a C3H female wild-type (WT) mouse. In the F1 generation, D/+ siblings were mated to generate homozygous (D/D) mice in the F2 generation. These mice were then monitored for seizure onset, lifespan and seizure frequency. This process was repeated for 28 times (N28) and evaluated the D/D mice each time to assess whether there were any differences in phenotype compared to D/D mice in B6 colony. In N6 (98.4%) line, we observed that D/D mice displayed three distinct phenotypes. In order to see this phenomenon was exclusive to N6 line, we re-performed all the backcrossing from new progenitors to test the possibility of recreating. While this effort did not re-produce the unusual long-lived phenotype, it resulted in the creation of the N6B line. We finished the second line of backcrossing with N7B line. When animals were euthanized, euthanasia was performed by cervical dislocation without anesthesia. No anesthetics were administered in this study. All animal work was performed in the animal facility in the Thomas Keating Building at the University of Arizona, Tucson, AZ, USA, and was approved by the University of Arizona Institutional Animal Care and Use Committee (IACUC protocol #16-160).

### 2.2 Genotyping

Tail biopsies were digested using 1 mg/mL proteinase K in a buffer containing 100 mM NaCl, 100 mM EDTA, 50 mM Tris-HCl (pH 8.0), and 1% SDS. DNA was subsequently extracted using phenol and chloroform, followed by ethanol precipitation. A 327 bp genomic region encompassing the targeted *Scn8a* site was amplified via PCR using primers Tar-F (5′-TGACTGCAGCTTGGACAAGGAGC-3′) and Tar-R (5′-TCGATGGTGTTGGGCTTGGGTAC-3′). The PCR products were digested with HincII and analyzed on 2% agarose gels to identify mice carrying the engineered HincII restriction site within the TALEN spacer region. The WT allele produces a single 327 bp fragment, whereas the mutant allele yields two fragments of 209 bp and 118 bp. For a complete description of the genotyping process, please refer to reference (11).

### 2.3 Phenotyping

A number of D/+ and D/D mice in 98.4% mouse line were monitored 24/7 using a video camera system to detect their first tonic-clonic seizure (TCS), record the number of TCS events during their lifespan, and determine the time of their death (12). For statistical analyses, age at first TCS and TCS per day in male D/D SL vs LL mice were compared with two-sided Mann–Whitney U tests after Shapiro–Wilk tests indicated non-normal distributions. Lifespan was compared between SL and LL using the log-rank test on Kaplan–Meier survival curves.

### 2.4 RNA-seq

Hippocampi samples from SL, LL, HP, and WT mice (n = 3 for each group) were harvested and flash frozen in liquid nitrogen and transferred to −80 °C freezer. The RNA-seq procedure was conducted at the University of Arizona Genetics Core. Briefly, the samples were processed for RNA extraction using 400-650 µl TRIzol reagent from Invitrogen (REF: 10296010) and PureLink™ RNA mini kit (Cat. No: 12183018A). Then Watchmaker mRNA library Prep Kit (Kit code: 7BK0001-384) was utilized to capture polyadenylated RNA transcripts, RNA fragmentation, cDNA synthesis, and library amplification with Equinox Master® Mix (Kit code: 7K0014-384). Paired-end FASTQ files for each hippocampal RNA-seq library were quality-checked with FastQC (per-read base quality, adapter content, and duplication). fastp then trimmed and filtered reads (adapter detection for paired-end libraries, quality trimming, and minimum read length), producing per-sample fastqc files and HTML/JSON QC reports.

Trimmed reads were aligned to the Mus musculus GRCm39 reference genome (NCBI Assembly GCF_000001635.27) with STAR, using a splice-aware index built from the same assembly and NCBI RefSeq gene annotation. Alignments were written as coordinate-sorted BAM files per sample. Gene-level read counts were obtained with featureCounts on STAR BAMs and the NCBI GRCm39 GFF, with paired-end counting. Counts were merged into a single matrix for downstream statistics.

For differential expression, we used edgeR in RStudio. Raw counts were loaded into a DGEList, lowly expressed genes were removed with filterByExpr, libraries were normalized with Trimmed Mean of M-value (log2 fold change) or TMM (calcNormFactors), and dispersions were estimated before pairwise exact tests of SL, LL, and HP versus WT (three biological replicates per group). Genes with false discovery rate (FDR) < 0.05 were treated as differentially expressed. For variant calling on the same STAR alignments, FreeBayes was run per sample against the GRCm39 reference (with samtools and bcftools for indexed BAMs and VCF handling) to produce per-sample VCFs for genome-wide variant screening. In order to see if the candidate variant (frameshift indel) found in whole-genome sequencing (WGS) exists in BAM files form RNA-seq data, first the T2T variant interval “15:113,714,890–113,714,892” was lifted to GRCm39 using UCSC liftOver against the published chain “GCA_964188535.1ToMm39.over.chain.gz” (chain source contig “OZ077256.1”); we verified the reference sequence at the lifted coordinates using samtools faidx against the GRCm39 reference genome. The *Scn8a*-N1768D variant was lifted over as expected to GRCm39 chr15:100,937,928 A>G, confirming the liftOver chain.

To examine the *Scn8a* frameshift variant candidate at read level, deletion-supporting (ALT; C) and reference (REF; CGT) reads were quantified at GRCm39 chr15:100,911,188 from hippocampal RNA-seq BAMs with a custom Python script. For each sample, primary alignments with MAPQ ≥ 10 were scanned over the three-base REF window; reads that did not fully cover the window, or that matched neither haplotype unambiguously, were excluded. Each spanning read was classified as REF, ALT, ambiguous, or other by exact sequence matching the REF or ALT haplotype. Informative depth was defined as REF + ALT reads per mouse. Per-mouse ALT fraction was ALT / (REF + ALT).

A dose-dependent increase in deletion-allele representation was tested with the Jonckheere– Terpstra (one-sided, increasing) ordered trend test on per-mouse ALT fractions, grouped by expected L-allele copy number at the indel (0: SL [D+/D+] and WT [++/++]; 1: LL [DL/D+]; 2: HP [DL/DL]). In SL and WT mice, non-L (REF) reads per genomic copy = REF ÷ 2. LL mice (DL/D+) carried one REF and one ALT allele, so non-L reads per copy = REF and L-bearing reads per copy = ALT. HP mice (DL/DL) carried two ALT alleles at the indel, so L-bearing reads per copy = ALT ÷ 2. Group means and Standard Error of Means (SEM) were calculated for each group. In LL heterozygotes, non-L read support per copy was compared with pooled SL and WT controls using a two-sided Mann–Whitney U test; L-bearing read support per copy was compared with controls using a one-sided Mann–Whitney U test (alternative: less). Heterozygous deletion-allele fractions in pooled long-read WGS DNA from dam (6389) and offspring (6527) were compared with pooled LL hippocampal RNA-seq by Fisher’s exact test. WGS trio pileup counts were obtained from WhatsHap haplotagged BAMs (see below).

### 2.5 Whole-Genome Long-Read Sequencing

Liver tissues from a long-lived heterozygous dam (D/+ LL), a short-lived heterozygous sire (SL D/+), and a long-lived homozygote (D/D LL) offspring were sent to Arizona Genetics Core at BIO5 for extraction of high-molecular-weight DNA. Then samples were sent to Arizona Genomics Institute for DNA library preparation and Pacific Biosciences (PacBio) long-read sequencing. High-molecular-weight (HMW) genomic DNA was extracted from each liver sample using the Nanobind PanDNA kit (PacBio, 103-260-000) according to the manufacturer’s animal-tissue protocol. For liver, 25–30 mg of tissue was finely minced on ice to ≤1 mm³ fragments, homogenized in cold Buffer CT with a TissueRuptor II (Qiagen), and lysed with Proteinase K and Buffer CLE3 at 55 °C with agitation (900 rpm, 30 min). After RNase A treatment and precipitation with Buffer SB, cleared lysate was bound to a Nanobind disk in the presence of isopropanol, washed sequentially with Buffers CW1 and CW2, and eluted in Buffer LTE. Eluates were homogenized by pipette mixing and allowed to solubilize overnight at room temperature before quality control.

DNA concentration and purity were assessed by triplicate NanoDrop One measurements (A260/A280 and A260/A230) and triplicate Qubit dsDNA Broad Range assays (Thermo Fisher Scientific) from the top, middle, and bottom of each eluate. HMW DNA size distribution was evaluated on a Femto Pulse system (Agilent Technologies) using the Genomic DNA 165 kb kit, with dilutions prepared using wide-bore pipette tips to minimize shearing.

For each sample, purified genomic DNA was sheared to 15–20 kb using a Megaruptor 3 (Diagenode) and purified with SMRTbell cleanup beads. SMRTbell libraries were constructed with the SMRTbell Prep Kit 3.0 (PacBio) following the manufacturer’s protocol. Libraries were size-selected on a Pippin HT (Sage Science), followed by a final SMRTbell cleanup bead purification. Finished libraries were quantified with the Qubit dsDNA HS Assay Kit (Invitrogen), and size distributions were confirmed on the Femto Pulse system (Agilent).

Libraries were prepared for sequencing using the Sample Setup and Run Design workflow in SMRT Link. Sequencing primer was annealed and polymerase-bound complexes were loaded onto 25M SMRT Cells. Sequencing was performed on a PacBio Revio instrument using Revio sequencing chemistry in circular consensus sequencing mode for 30 h per run, targeting approximately 30× genome coverage per sample.

HiFi reads were aligned to the Mus musculus B6 telomere-to-telomere (T2T) reference genome (NCBI Assembly GCA_964188535.1) with pbmm2, and alignments were sorted and indexed with samtools. Single-nucleotide variants (SNVs) and small insertions/deletions (indels) were called jointly for the trio with bcftools, producing one multi-sample VCF restricted to chr15:90,000,000–116,793,490 on the T2T B6 assembly because the locus of interest is tightly linked to *Scn8a*-N1768D on chromosome 15. To resolve parental origin of alleles, the merged SNV/indel VCF and aligned BAM files were phased as a dam–sire–offspring trio with WhatsHap, and haplotype (Ht) and phase-set (PS) tags were written to the BAMs with whatshap haplotag.

Candidates for the variant from trio-combined VCF were enumerated against four rules: 1. The variant must be within ±3 Mb of *Scn8a*-N1768D; 2. offspring phased heterozygous with ALT on the maternal haplotype; 3. dam phased with ALT on her *Scn8a*-N1768D-carrying haplotype, or homozygous-ALT; 4. sire phased REF on his *Scn8a*-N1768D-carrying haplotype, or homozygous-REF. The candidate table was annotated against the T2T GFF3 for gene identity, biotype, and location class (coding, 5′UTR, 3′UTR, exonic but non-coding, intronic, promoter, intergenic, or mixed); the GRCm39 ortholog projection was used only as a name source when a T2T-only Ensemble gene ID could not be resolved by “www.mygene.info”.

To find the origin of the candidate variant found in WGS, we mapped the T2T variant coordinate onto the C3H assembly (GCA_921997125.2; chr15 contig OW971850.1) using minimap2. First, we aligned a 5,001-bp T2T window centered on the variant (chr15:113,712,390–113,717,390) to C3H chr15 and inferred the orthologous base at single-nucleotide resolution from the colinear alignment. To exclude a window-edge artifact, we repeated the analysis with an independent whole-chromosome alignment of C3H chr15 (101.3 Mb) to T2T chr15 (116.8 Mb), again with minimap2.

## 3. Results

### 3.1 Phenotypic Variability in D/+ and D/D Mice with Mixed C3H and B6 Genetic Backgrounds

D/D mice in the 98.4% colony showed marked phenotypic heterogeneity in age at death. Among 236 D/D mice with known age at death or euthanasia, three phenotypes were distinguished. The first was HP (n = 48; 20.3%), with a mean age at euthanasia of 16.7 days (SD, 2.3 days). In these mice, hindlimb paralysis became discernible around P14, progressed daily, and by P21 most could not move or eat (**Video S1**). They were humanely euthanized before their natural death to prevent suffering, and their lifespan data were excluded from survival analysis. A second phenotype was SL (n = 63; 26.7%), with a mean lifespan of 20.9 days (SD, 1.8 days) (**Figure 1A**). Like SL mice on the B6 background, these animals typically died around P21 after several back-to-back TCS. The third group comprised mice surviving beyond 25 days, classified as LL (n = 125; 53.0%), with a mean lifespan of 64.5 days (SD, 22.5 days), reflecting substantial variability in survival among LL mice (**Figure 1A**). Colony-wide proportions and mean lifespans are summarized in **Table 1**; the individually monitored cohort is detailed per animal in **Table S1**.

**Figure 1.**
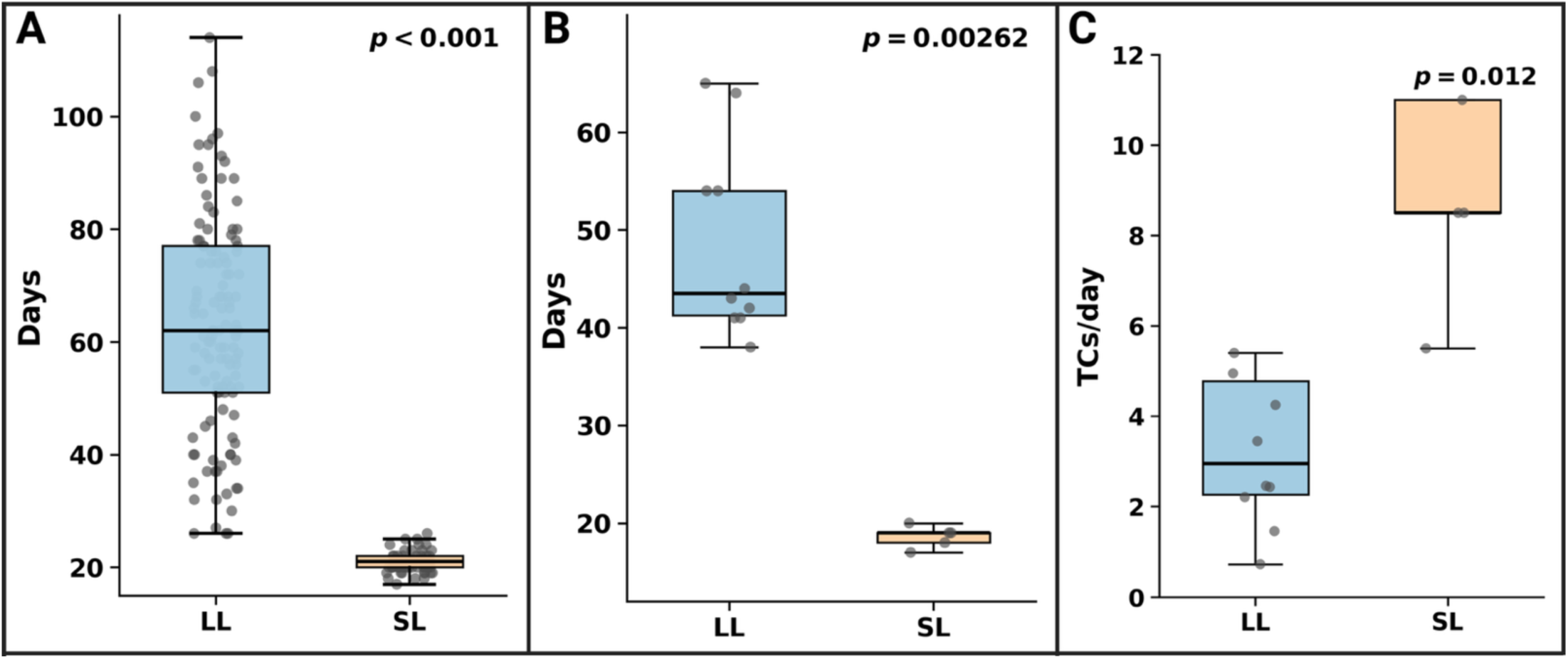
Seizure characteristics in LL vers us SL mice. (males; seizure metrics were available only for male SL mice). **A**) Lifespan, **B**) Age at seizure onset. **C**) seizures per active day. boxes show median and IQR, points are individual mice, p-values from Mann–Whitney U tests. Summary statistics for each metric are presented in **Table 1**. Created in BioRender. Bahramnejad, E. (2026) https://BioRender.com/es108an.

**Table 1.** Survival and seizure outcomes in male *Scn8a*-N1768D mice on the 98.4% C3H.B6 background.

| Genotype | Sex | Variable | Unit | n<br>(HP) | Mean ±<br>SD<br>(HP) | n<br>(SL) | Mean ±<br>SD (SL) | n<br>(LL) | Mean ±<br>SD (LL) | p-value<br>(SL vs<br>LL) |
| --- | --- | --- | --- | --- | --- | --- | --- | --- | --- | --- |
| D/D | Male | Lifespan<br>or Age at<br>euthanasia | Days | 48 | 16.7 ±<br>2.3 | 63 | 20.9 ±<br>1.8 | 125 | 64.5 ±<br>22.5 | <0.0001 |
| D/D | Male | TCS onset | Days | - | - | 5 | 18.6 ±<br>1.1 | 10 | 48.6 ±<br>9.9 | 0.003 |
| D/D | Male | TCS<br>frequency | TCS<br>per<br>day | - | - | 5 | 13.5 ±<br>11.6 | 10 | 5.6 ± 8.2 | 0.012 |
| D/+ | Male | Lifespan<br>or Age at<br>euthanasia | Days | - | - | 29 | 108.8 ±<br>27.0 | 10 | 218.9 ±<br>194.4 | 0.038 |
| D/+ | Male | TCS onset | Days | - | - | - | - | 5 | Not<br>observed | - |

### 3.2 Age of Onset

Mann–Whitney U tests were used to compare age at seizure onset between male SL and male LL D/D mice, as a Shapiro–Wilk test indicated non-normality in the LL group (p = 0.047). The pairwise comparisons revealed significant phenotype effects on TCS onset timing between SL and LL mice. LL males developed seizures significantly later than SL males (mean: 48.6 vs 18.6 days, p = 0.003), representing a meaningful delay of approximately 30 days in average (**Figure 1B**). These findings demonstrate that the LL phenotype is associated with delayed TCS initiation in male mice. Per-animal onset values are given in **Table S2** and summarized by phenotype in **Table 1**.

### 3.3 Number of Tonic-Clonic Seizures per Day

Mann–Whitney U tests were used to compare seizure frequency between male SL and male LL D/D mice, as Shapiro-Wilk normality tests revealed non-normal distributions across all groups for this variable (all p < 0.05). The analysis revealed significant phenotype effects on seizure frequency in males, with LL males experiencing significantly fewer TCS per day compared to SL males (mean: 5.6 vs 13.5 TCS/day, p = 0.012), representing an approximately 2.5-fold reduction in seizure burden (**Figure 1C**). This suggests that LL male mice had reduced seizure burden. Per-animal seizure-frequency values are given in **Table S2** and summarized in **Table 1**.

### 3.4 Kaplan-Meier Survival Curves

#### Homozygotes

To compare SL and LL survival directly in males, we performed Kaplan–Meier analysis on 188 male D/D mice with assigned SL or LL phenotype (63 SL and 125 LL), excluding HP mice euthanized for humane reasons (**Figure 2A**). Male SL survival fell sharply between approximately P18 and P26, whereas male LL survival declined much more gradually thereafter. A log-rank test showed a highly significant difference between SL and LL (p < 0.0001). (Per-animal survival in **Table S1**).

**Figure 2.**
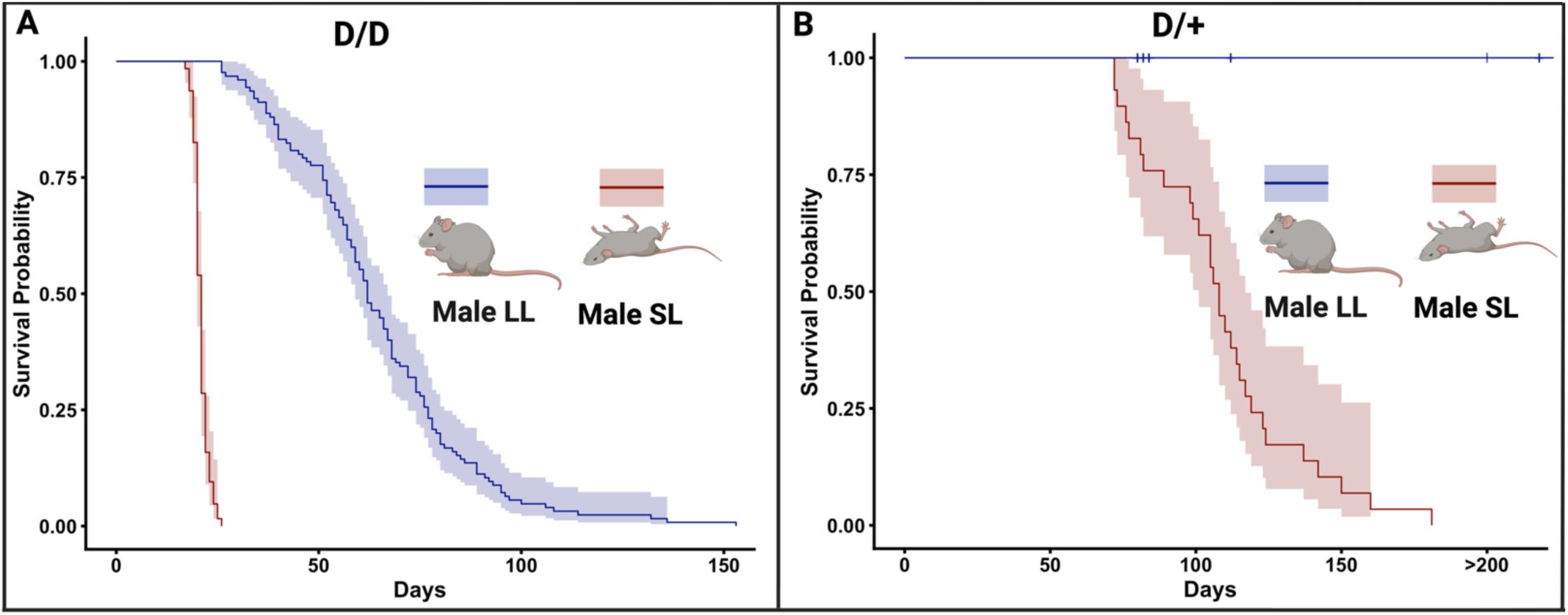
Survival of homozygous (D/D) and heterozygous (D/+) mice. **A**) Kaplan–Meier survival of D/D male mice with an assigned SL or LL phenotype. **B**) Kaplan–Meier survival of D/+ males with an assigned SL or LL phenotype Shaded areas represent 95% confidence intervals. Tick marks denote censored animals. Created in BioRender. Bahramnejad, E. (2026) https://BioRender.com/0by0v25.

#### Heterozygotes

As in homozygotes, heterozygous *Scn8a*-N1768D mice on the 98.4% background manifested two distinct phenotypes. In the first group, mice developed epilepsy and died during monitoring or were found dead in their cages—a phenotype closely resembling D/+ mice on the B6 background. We classified these animals as short-lived (D/+ SL). In the second group, mice did not develop epilepsy, survived the monitoring period without epilepsy-related death, and were euthanized at different ages. We classified these animals as long-lived (D/+ LL). Breeding outcomes were consistent with these phenotypes: D/+ SL × D/+ SL crosses produced SL D/D offspring; D/+ LL × D/+ SL crosses yielded LL D/D progeny; and D/+ LL × D/+ LL crosses yielded HP homozygotes.

Survival analysis of male D/+ mice on the 98.4% background used time to epilepsy-related death, with D/+ LL animals right-censored at euthanasia and D/+ SL animals scored as events when found dead (**Figure 2B**). Survival differed strongly between D/+ LL and D/+ SL groups (log-rank χ² = 20.2, p = 6.9 × 10⁻⁶). D/+ SL males had a mean age at death of 108.8 days (SD, 27.0; n = 29). Among D/+ LL males, no epilepsy-related deaths occurred; these animals were euthanized at different ages rather than reaching the death endpoint (n = 10). D/+ LL males had markedly longer survival without epilepsy-related death than D/+ SL males. Overall, survival among male D/+ mice separated sharply according to phenotype. (Per-animal survival in **Table S1**).

### 3.5 Susceptibility to Audiogenic Seizures

Wengert et al. reported that *Scn8a*-N1768D mice are susceptible to audiogenic seizures and that jingling keys (key test) can elicit TCS in D/+ animals (13). Guided by those findings, we tested D/+ SL and D/+ LL mice (n = 10 per group) for susceptibility to audiogenic TCS. D/+ SL mice responded to the stimulus with a TCS; D/+ LL mice did not (**Video S2**), resembling WT animals, which likewise do not respond to the key test (Fisher’s exact test, two-sided p = 1.1 × 10⁻⁵). During euthanasia by cervical dislocation, every D/D and D/+ SL mouse exhibited a tonic seizure immediately before death, whereas none of the D/+ LL mice did, resembling WT animals. Together with the key test, these observations provided additional evidence that D/+ LL mice lacked the epileptic signs and symptoms seen in D/+ SL mice.

### 3.6 Two-Locus Model of Inheritance

Genetic analysis of our 98.4% colony identified a second locus “L” on chromosome 15 tightly linked to *Scn8a*-N1768D, with genotypes segregating for homozygotes as D+/D+ (SL), DL/D+ (LL), and DL/DL (HP) and heterozygotes as D+/++ for D/+ SL and DL/++ for D/+ LL. Two independent lines of evidence support this two-locus model and the tight linkage between the second locus and *Scn8a*-N1768D: (1) only five recombinants were detected among 266 litters, corresponding to an estimated genetic distance of ∼1.88 cM, and (2) each litter from D/+ x D/+ cross consistently produced a single-phenotype (SL, LL, or HP) D/D mice, demonstrating that the “L” locus and *Scn8a*-N1768D co-segregate as a linked unit.

We quantified the strength of linkage using standard Logarithm of the Odds (LOD) analysis based on the observed recombinant fraction. With 5 recombinants among 266 litters, the estimated recombination fraction is θ̂ = 0.0188 (≈1.88 cM; 95% CI: 0.00613–0.0433). The LOD curve peaks at θ̂ ≈ 0.0187 with LOD(max) = 69.29, providing overwhelming statistical support for tight linkage between the “L” locus and *Scn8a*-N1768D (**Figure 3A**). For context, a LOD score above 3.0 is considered significant evidence for linkage; our LOD score of 69.29 exceeds this threshold by more than 20-fold, establishing unequivocal genetic linkage.

**Figure 3.**
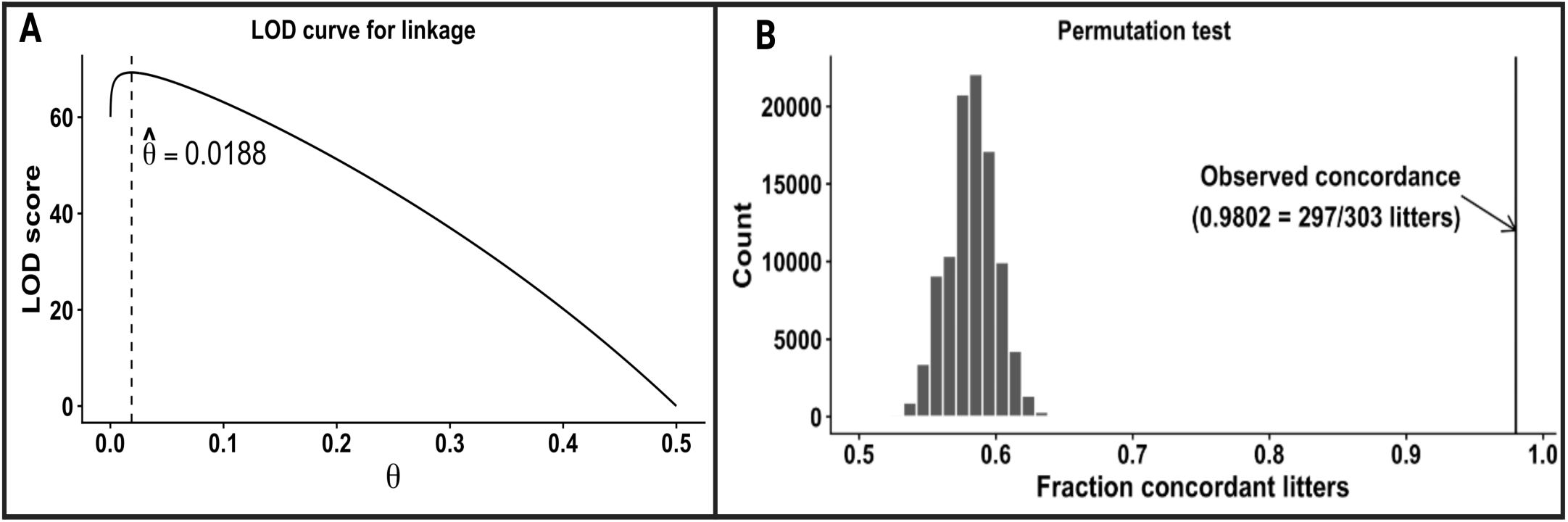
Genetic evidence for tight linkage between the L modifier and *Scn8a* on chromosome 15. **(A)** LOD score plotted as a function of recombination fraction θ from 5 recombinants among 266 informative litters. The curve compares the likelihood of linkage at θ to no linkage (θ = 0.5). The dashed line marks the maximum-likelihood estimate θ̂ = 0.0188. **(B)** Permutation test for litter phenotype concordance among D/D mice. The histogram shows the null distribution of the fraction of concordant litters obtained by randomly shuffling phenotype labels across pups while preserving litter sizes and overall SL/LL/HP frequencies. The solid line marks the observed concordance (297/303 litters; 0.98). Created in BioRender. Bahramnejad, E. (2026) https://BioRender.com/7giovdb.

Our results show that in each litter from D/+ × D/+ crosses, D/D mice only showed one of the three phenotypes; they were either SL, LL, or HP, and only when we crossed a D/D × D/+ or D/D, we would see more than one phenotype in the D/D progeny. We assessed the statistical significance of the “single phenotype per litter” pattern for D/D using a permutation test that preserves litter sizes and overall SL/LL/HP frequencies while randomly shuffling phenotypes across pups to generate a null distribution under the hypothesis of independent segregation (**Figure 3B**). In the observed data, 297/303 litters (98.0%) were concordant (single phenotype per litter). In those 6 litters that we observed more than one phenotype among D/D mice, we crossed D/D × D/+ or D/D. The observed concordance lies far outside the null distribution (p = 1×10⁻⁵, 100,000 permutations), demonstrating that the litter-level consistency does not arise by chance and confirming co-segregation of the “L” locus with *Scn8a*-N1768D. This extremely low p-value indicates that the probability of observing this degree of litter concordance by chance alone is less than 1 in 100,000, providing robust statistical validation that the “L” locus and *Scn8a*-N1768D are genetically linked.

### 3.7 LL and HP Mice Show Significant Reduction in the *Scn8a* mRNA Levels

In line with the two-locus model, SL, LL, and HP mice are all *Scn8a*-N1768D homozygotes but differ at the linked “L” locus: SL is D+/D+ (no L allele), LL is DL/D+ (one L allele), and HP is DL/DL (two L alleles). Hippocampal RNA-seq results showed that relative to WT, SL showed no significant change in *Scn8a* expression (edgeR log₂ fold change = −0.10; FDR = 0.51), whereas LL showed ∼30% lower *Scn8a* mRNA (log₂ fold change = −0.57; FDR = 0.009) and HP showed ∼70% lower *Scn8a* mRNA (log₂ fold change = −1.75; FDR = 3.1 × 10⁻⁶). Mean hippocampal *Scn8a* expression was 8.44 log₂(CPM) in WT, 8.34 in SL, 7.88 in LL, and 6.68 in HP (**Figure 4A**). This graded, statistically significant reduction supports a dose model in which SL carries the highest mutant-channel transcript level, LL an intermediate level, and HP the lowest, and it provides a mechanistic axis for interpreting downstream transcriptional differences among phenotypes.

**Figure 4.**
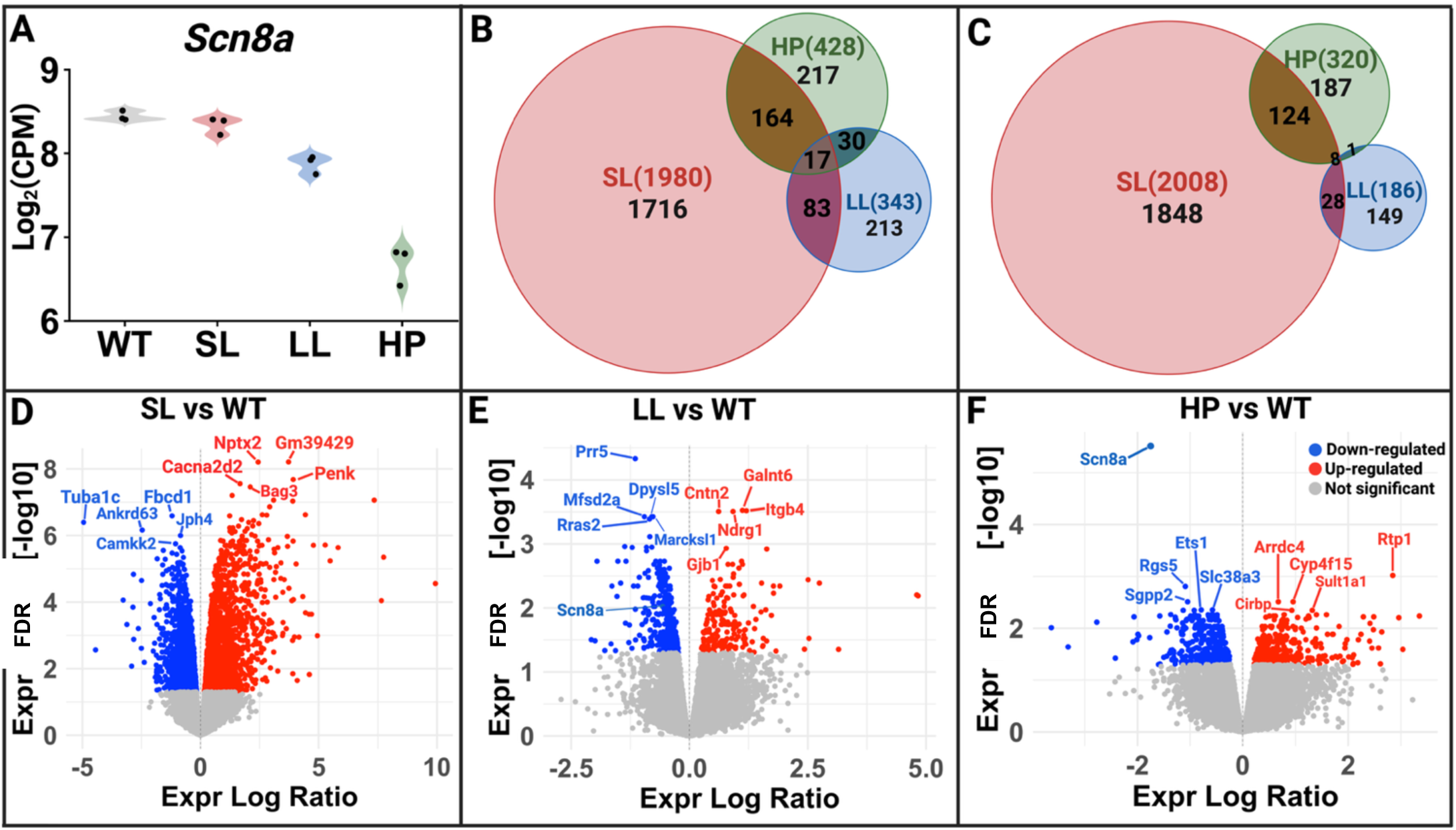
Differential gene expression across three phenotypes in D/D mice (FDR < 0.05). **(A)** Violin plot of *Scn8a* expression in hippocampus of WT, SL, LL, and HP males. **(B)** Venn diagram of downregulated differentially expressed genes (DEGs) in SL, LL, and HP versus WT. **(C)** Venn diagram of upregulated DEGs in SL, LL, and HP versus WT. **(D–F)** Volcano plots of DEGs in SL, LL, and HP versus WT, respectively. The top five downregulated and top five upregulated genes by FDR are labeled in each volcano plot. In volcano plots, red indicates upregulated genes, blue indicates downregulated genes, and grey indicates genes that are not significant. Created in BioRender. Bahramnejad, E. (2026) https://BioRender.com/mvr6qgl.

### 3.8 Phenotype-Specific Differential Expression

To quantify overlap among phenotypes, we compared hippocampal differential expression in SL, LL, and HP versus WT mice (FDR < 0.05) and summarized shared up- and downregulated genes in separate Venn diagrams (**Figure 4B-C**). Downregulated genes followed the same pattern at larger scale (1,980 in SL, 343 in LL, and 428 in HP), with 1,716 SL-specific, 213 LL-specific, and 217 HP-specific genes. SL and HP shared 181 downregulated genes, SL and LL shared 100, and LL and HP shared 47; 17 genes were downregulated in all three (**Figure 4B**). Upregulated genes numbered 2,008 in SL, 186 in LL, and 320 in HP; most SL genes were phenotype-specific (1,848), whereas 149 and 187 were unique to LL and HP. Pairwise overlap was greatest between SL and HP (132 genes), compared with 36 between SL and LL and 9 between LL and HP; only 8 genes were upregulated in all three genotypes (**Figure 4C**). In both directions, LL had the fewest differentially expressed genes and SL the most, with HP intermediate—a pattern compatible with the moderate phenotype of LL mice relative to SL and HP. Together, these overlaps indicate that most differentially expressed genes were significant only in SL, whereas multi-phenotype overlap was greatest between SL and HP and comparatively limited between LL and HP.

We next displayed genome-wide differential expression in volcano plots for each phenotype versus WT (**Figure 4D–F**). In SL versus WT, top downregulated genes included *Fibcd1*, *Tuba1c*, *Ankrd63*, *Jph4*, and *Camkk2*, and top upregulated genes included *Gm39429*, *Nptx2*, *Penk*, *Cacna2d2*, and *Bag3* (**Figure 4D**). LL versus WT yielded a smaller set of high-confidence changes, with *Prr5*, *Dpysl5*, *Mfsd2a*, *Marcksl1*, and *Rras2* among the top downregulated genes and *Galnt6*, *Itgb4*, *Ndrg1*, *Cntn2*, and *Gjb1* among the top upregulated genes (**Figure 4E**). HP versus WT was distinguished by *Scn8a* as the most significantly downregulated gene; other top downregulated genes included *Rgs5*, *Sgpp2*, *Ets1*, and *Slc38a3*, and top upregulated genes included *Rtp1*, *Arrdc4*, *Cyp4f15*, *Sult1a1*, and *Cirbp* (**Figure 4F**). Pathway-level interpretation of these phenotype-specific programs is presented in the companion paper (Hammer and Bahramnejad, in prep).

### 3.9 Rationale for Long-Read Trio Whole-Genome Sequencing

Following differential gene-expression analysis, we searched the 12-mouse RNA-seq VCFs for a second-locus variant consistent with our two-locus model. Linkage mapping placed this locus within 1.88 cM of *Scn8a*-N1768D; we therefore searched ±3 Mb around N1768D to avoid missing a marginally linked variant. Filtering per-sample VCFs for cohort-matching genotypes (WT and SL, REF/REF; LL, ALT/REF; HP, ALT/ALT) yielded a single candidate: a GTG>ACA substitution in *Spryd3* intron 9 (GRCm39 chr15:102,033,991). Because an intronic *Spryd3* variant is unlikely to modulate *Scn8a* transcript level, we re-screened the linked region at base-pair resolution by PacBio HiFi WGS of a trio comprising a D/+ LL dam (DL/++), a D/+ SL sire (D+/++), and their LL D/D offspring (DL/D+).

### 3.10 Candidate identification across WGS and RNA-seq data and its origin

VCF candidates passing the four rules described in Methods for the WhatsHap-phased trio VCF are listed in **Table S3** (n = 4,005). Each variant was annotated against the T2T GFF3 for gene identity, biotype, and genomic context (coding, 5′ UTR, 3′ UTR, intronic, promoter, intergenic, mixed, or exonic non-coding). Most candidates were intronic or intergenic: 1,880 (46.9%) and 1,658 (41.4%), respectively; 59 (1.5%) were coding, 236 (5.9%) promoter, 137 (3.4%) 3′ UTR, 28 (0.7%) 5′ UTR, 6 (0.1%) mixed, and 1 (<0.1%) exonic non-coding. The set comprised 2,261 SNVs and 1,744 indels.

Twelve candidates mapped to *Scn8a* (11 intronic, 1 coding). The sole coding variant was a 2-bp deletion at T2T chr15:113,714,890 (CGT→C), 26,744 bp downstream of N1768D. Overall, the WhatsHap VCF-only screen yielded thousands of *cis*-linked candidates enriched for non-coding variation, with a small coding subset that included this single *Scn8a* frameshift indel.

**Figure 5** summarizes phased haplotype structure at the frameshift and N1768D loci in the WGS trio. Given the dam’s D/+ LL and the sire’s D/+ SL phenotypes, the frameshift variant is expected to lie in *cis* with N1768D on the dam’s haplotype and to be WT on the sire’s N1768D-carrying haplotype; in the LL offspring, it should therefore appear only on the maternal haplotype. As shown in **Figure 5**, the phased frameshift indel matches this expectation. Because observed phasing agrees with the configuration predicted from parental phenotypes, the variant is supported as a candidate for “L” allele.

**Figure 5.**
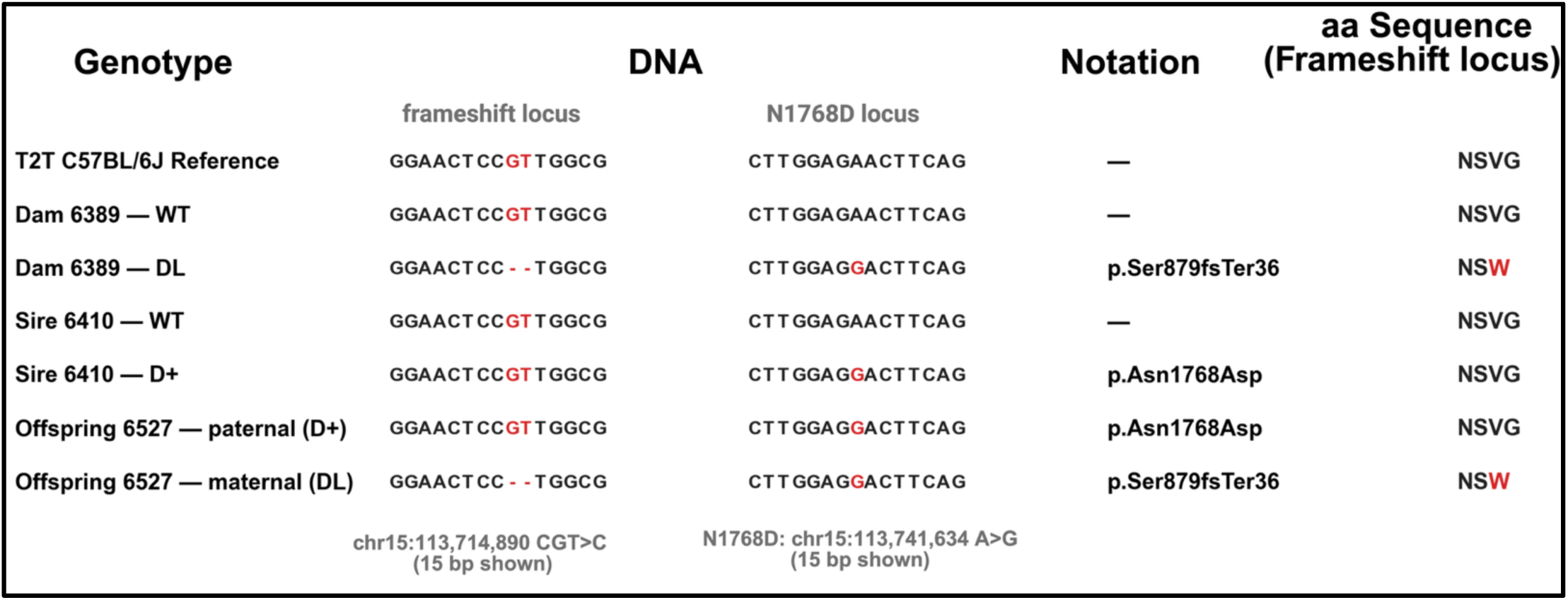
Phased *Scn8a* haplotypes at the frameshift (chr15:113,714,890 CGT>C) and N1768D (chr15:113,741,634 A>G) loci in the WGS trio. Fifteen-base DNA windows, Human Genome Variation Society protein notation, and local amino-acid sequence at the frameshift site are shown for the T2T C57BL/6J reference, dam 6389 (++ and DL), sire 6410 (++ and D+), and offspring 6527 (paternal D+, maternal DL). The frameshift deletion (p.Ser879fsTer36) is carried in *cis* with N1768D on the maternal haplotype only. Created in BioRender. Bahramnejad, E. (2026) https://BioRender.com/4x8b916.

We next asked whether this coding indel is detectable in bulk RNA-seq from the 12-mouse hippocampal cohort (n = 3 per WT, SL, LL, and HP). At the lifted GRCm39 coordinate, per-sample VCFs called the deletion only in HP mice (homozygous ALT) and not in WT, SL, or LL— a known limitation of short-read indel calling at moderate depth, especially for heterozygous indels (14). Read-level REF and ALT counts from each STAR-aligned BAM files (Methods; **Table S4**) showed zero ALT reads in all WT and SL replicates (ref/ref), zero REF reads in all HP replicates (del/del), and both REF and ALT reads in all LL replicates (ref/del). All 12 animals matched the genotype expected from the two-locus model. ALT fraction increased with expected deletion dosage (SL and WT 0 → LL 1 → HP 2 copies), and the Jonckheere–Terpstra test supported this increasing trend (z = 3.38, one-sided p = 3.6 × 10⁻⁴). This dose pattern is consistent with the dose-dependent reduction of *Scn8a* mRNA reported above. We therefore retain GRCm39 chr15:100,911,188–100,911,190 (REF CGT, ALT C) as the candidate “L” allele in *cis* with *Scn8a*-N1768D, supported by evidence in both the WGS trio and the RNA-seq cohort.

Finally, we asked whether the frameshift variant was inherited from the C3H background rather than arising *de novo*. Alignment of a 5,001-bp T2T window containing the variant to the C3H genome (OW971850.1) placed T2T chr15:113,714,890 at OW971850.1:98,185,433 with >99% identity, and flanking sequence was identical between T2T B6 and C3H; both references carry REF “CGT” at the orthologous site. The deletion is therefore unlikely to have been inherited from C3H strain and instead arose *de novo* in the colony. Together, phased trio WGS, read-level RNA-seq validation, and strain-reference comparison identify a single *Scn8a* coding frameshift in *cis* with N1768D as the “L” allele and support its *de novo* origin in the congenic line.

### 3.11 Predicted Molecular Consequence of the L-allele Indel

The deletion lies within coding exon 15 of the canonical *Scn8a* transcript (*ENSMUST00365135273*; 26 exons in the T2T B6 annotation), removing two nucleotides immediately after codon 879 of the encoded Na_V_1.6 channel and shifting the downstream reading frame. Translation of the mutant allele terminates at a premature termination codon (PTC) at codon 914 (TAA). This termination codon lies 26,131 nucleotides upstream of the 3′-most exon–exon junction—far beyond the >50-nucleotide threshold for nonsense-mediated mRNA decay (NMD) (15, 16). The resulting truncation falls between domain II and domain III of Na_V_1.6 and would eliminate the DIII/DIV transmembrane core, the DIII–DIV inactivation linker, and the C-terminal cytoplasmic tail—including the N1768D residue itself. Any L-bearing *Scn8a* mRNA that escapes such clearance would still encode only the truncated, most likely non-functional channel architecture outlined in **Video S3**.

Since we observed dose-dependent reduction of total *Scn8a* mRNA in LL and HP mice carrying one and two copies of the “L” indel, respectively, but not in SL mice carrying N1768D without “L” (**Figure 4**), we next asked whether that loss reflects selective reduction of the indel-bearing transcript rather than uniform down-regulation of all *Scn8a* mRNA. To test this, we counted reads only at the 2-bp frameshift indel. If the “L” transcript is selectively unstable, heterozygotes at the indel site should still carry the deletion on one of two *Scn8a* chromosomes in genomic DNA, so read counts at the indel in DNA should be ∼50% REF and ∼50% ALT (Mendelian heterozygosity), whereas read counts at the same indel in mRNA should show far fewer ALT than REF reads if the L-bearing transcript is selectively lost. Total *Scn8a* mRNA should then decline as “L” copy number increases even though N1768D is present in all mutant phenotypes. We tested this in hippocampal RNA-seq from 12 mice (++/++, D+/D+, DL/D+, and DL/DL) and, for genomic reference at the same indel coordinate, in long-read WGS DNA from the two trio members heterozygous at the indel—dam 6389 (DL/++) and offspring 6527 (DL/D+) (**Figure 6**).

**Figure 6.**
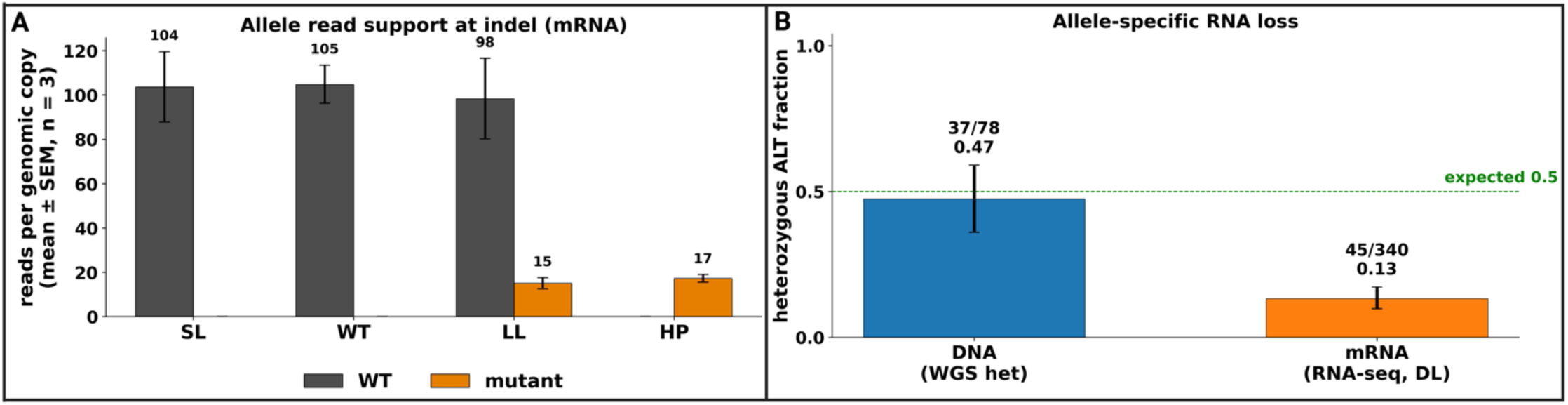
Allele-specific read support at the *Scn8a* frameshift indel. **(A)** REF (non-L) and ALT (L-bearing) informative reads per genomic copy at chr15:100,911,188 in hippocampal RNA-seq (n = 3 per group: SL, WT, LL, HP). **(B)** Heterozygous deletion-allele fraction at the indel in pooled WGS DNA (dam 6389 + offspring 6527) versus pooled LL RNAseq; dashed line, Mendelian heterozygote expectation (0.5). Created in BioRender. Bahramnejad, E. (2026) https://BioRender.com/pigkfp4.

At the indel, spanning reads were classified as REF or ALT by exact match and normalized per genomic copy (**Figure 6A**). SL and WT showed only non-L reads (∼104–105/copy); LL showed preserved non-L output (∼98/copy) but strongly reduced L-bearing reads (∼15/copy); HP showed only L-bearing reads (∼17/copy), as expected for DL/DL genotype but at low absolute depth. Non-L support in LL did not differ from ref/ref controls (two-sided Mann–Whitney U, p = 0.905), whereas L-bearing support was reduced (one-sided Mann–Whitney U, p = 0.012). Loss is therefore allele-specific and *cis*-acting on the DL transcript, not uniform down-regulation of both alleles.

Comparing heterozygous mice directly, deletion-allele fraction at the indel was near Mendelian expectation in DNA but markedly reduced in mRNA (**Figure 6B**). Pooled WGS DNA from dam and offspring at the indel was 0.47 (37/78 ALT reads), close to the expected 0.5, whereas pooled LL hippocampal RNA-seq at the same site was 0.13 (45/340; Fisher’s exact test, p = 2.6 × 10⁻¹⁰). The RNA deficit in LL mice is therefore not explained by loss of the deletion from the genome, but by selective under-representation of L-bearing reads in the transcript pool. In summary, read-level analysis at the frameshift indel shows selective, *cis*-acting loss of L-bearing transcripts with preserved non-L read support and intact genomic dosage, yielding dose-dependent reduction of total *Scn8a* mRNA rather than uniform down-regulation of both alleles.

## 4. Discussion

*SCN8A* encodes the voltage-gated sodium channel Na_V_1.6, a principal driver of neuronal excitability in the central nervous system (17). In humans, *de novo SCN8A* variants produce a clinical spectrum that tracks their functional effect: GoF variants cause DEE with early-onset, frequently drug-resistant seizures, whereas LoF variants cause intellectual disability or milder generalized epilepsy without the severe seizure burden (18–21). A striking feature of these disorders is the marked phenotypic variability among individuals carrying the same pathogenic variant, implicating genetic modifiers in disease severity and course (22). The *Scn8a*-N1768D knock-in mouse reproduces the GoF arm of this spectrum—heterozygotes develop TCS and sudden unexpected death in epilepsy (23)—providing a tractable system in which such modifiers can be found. Because modifiers can expose both disease mechanism and therapeutic entry points, and because *SCN8A* behaves as a “Goldilocks” gene in which both excess and deficit of Na_V_1.6 are deleterious, a modifier that re-tunes Na_V_1.6 dose is of particular translational interest (24).

Here we identify the genetic modifier that converts the uniformly severe *Scn8a*-N1768D GoF phenotype into a graded series of outcomes, and show that it is a *de novo*, *cis*-acting variant within *Scn8a* itself. Phenotyping defined three homozygous classes (SL, LL, HP) and two heterozygous classes (D/+ SL, D/+ LL) that form a dose-ordered series; linkage placed a single modifier locus in tight linkage with *Scn8a* (LOD 69.3; ∼1.88 cM) that co-segregates with phenotype litter-by-litter (Linkage); and *Scn8a* mRNA declined in proportion to modifier copy number (∼0%, ∼30%, ∼70%; mRNA dose). Long-read trio DNA sequencing resolved this locus to a 2-bp frameshift deletion in coding exon 15 of *Scn8a*, *de novo* and predicted to trigger NMD. Below we discuss that an intragenic, *de nov*o modifier of a disease gene is itself notable (4.1); that heterozygous *SCN8A* frameshifts span a broad human phenotypic spectrum, whereas in our colony one copy of “L” on the N1768D allele converts D/+ SL into a TCS-resistant D/+ LL state (4.2); that copy number of this LoF allele titrates Na_V_1.6 through a narrow “Goldilocks” window in which both excess and near-complete loss are deleterious (4.3); that this frameshift mutation that this genetic rescue validates *Scn8a*-lowering as a therapeutic strategy while defining the window such a therapy must respect (4.4); that the GoF and LoF poles of one gene therefore demand opposite therapeutic logic (4.5); and that the resulting graded molecular axis links genotype to the phenotype-specific transcriptional programs dissected in the companion paper (4.6).

### 4.1 A *De Novo*, Intragenic Modifier of *Scn8a*

Most known modifiers of monogenic epilepsies are trans-acting genes identified through strain-dependent differences in severity. In the *Scn1a*+/− model of Dravet syndrome, for example, survival is governed by modifier loci on several chromosomes (Dsm1–Dsm5), and the Dsm1 modifier was ultimately resolved to a regulatory variant of *Gabra2*, a GABA_A_-receptor subunit gene located on a different chromosome from the primary disease gene (25, 26). The modifier we describe differs on two fundamental counts: it lies within *Scn8a* itself, in *cis* with the N1768D pathogenic variant, and it arose *de novo* rather than as an inherited strain polymorphism. To our knowledge this is the first description of a *de novo*, intragenic, *cis*-acting modifier that re-tunes the expression of the very gene carrying the primary mutation. Because the modifier and the pathogenic variant reside on the same allele, their effects are inseparably linked at transmission, which both explains the clean litter-by-litter co-segregation we observe and distinguishes this architecture from classical trans-modifier genetics.

### 4.2 Impact of Heterozygous Frameshift Mutations in Human and Mouse *SCN8A*/*Scn8a*

In contrast to D/+ SL mice, male D/+ LL mice lacked spontaneous TCS and epilepsy-related death, were resistant to audiogenic TCS, and did not display a terminal tonic seizure at euthanasia by cervical dislocation, resembling WT mice. Because electroencephalogram, behavioral, and social testing were not performed, absence seizures and non-seizure phenotypes cannot be ruled out among D/+ LL mice.

Human heterozygous *SCN8A* frameshift variants show a broad phenotypic spectrum: intellectual disability, ataxia, and cerebellar atrophy without epilepsy (p.Pro1719ArgfsTer6); intermediate epilepsy with absence seizures (p.Asn544fsTer39); mild intellectual disability without epilepsy (p.Pro577ArgfsTer61); later-onset TCS with moderate intellectual disability (p.Met412IlefsTer47); early-onset DEE (p.Met1481IlefsTer12); and unclassifiable epilepsy with severe intellectual disability (p.Ala1622IlefsTer64) (18, 27, 28). Hack et al. (2023) reported nine frameshift cases, with phenotypes ranging from neurodevelopmental disorder without epilepsy to generalized epilepsy and DEE; some cases were unclassifiable or incompletely phenotyped (29).

In mice, heterozygous *Scn8a* frameshift alleles are associated with seizure resistance without a severe motor phenotype. Inglis et al. (2020) showed that Δ35/+ frameshift heterozygotes had no motor deficits, a normal acoustic startle response, and resistance to induced seizures (30). Consistent with a protective effect of inactivating a GoF allele, allele-specific CRISPR editing that introduces frameshifting indels in *cis* on *Scn8a*-N1768D rescues seizures and lethality in D/+ mice (31). Our D/+ LL mice lack spontaneous and audiogenic TCS and epilepsy-related death, paralleling the protective effect of CRISPR-mediated frameshift inactivation of the pathogenic allele in D/+ mice (31). Behavioral testing of D/+ LL versus WT mice is still needed to evaluate non-seizure outcomes.

### 4.3 Dose-Dependent Na_V_1.6 and the Goldilocks Window

Copy number of the LoF L allele titrates *Scn8a* mRNA, and therefore Na_V_1.6, in graded steps: no reduction in SL (D+/D+), ∼30% in LL (DL/D+), and ∼70% in HP (DL/DL). These graded reductions reflect *Scn8a* mRNA levels and do not necessarily represent Na_V_1.6 protein abundance; a similar dissociation between mRNA and protein levels has been reported for *SCN2A* (32). The phenotypic consequence is non-monotonic. Full GoF (SL) is lethal through severe early seizures, near-complete loss (HP) is lethal through hindlimb paralysis, and only the intermediate reduction (LL) rescues. This single allelic series recapitulates the opposite poles of human *SCN8A* disease, in which GoF variants cause focal DEE while LoF variants cause generalized epilepsy or intellectual disability without seizures (18, 19, 33). The HP phenotype closely mirrors the murine *Scn8a*-null (med) allele, which causes HP and juvenile lethality, rather than the surviving, ataxic phenotype of hypomorphic alleles such as med-J (17, 34).

Future studies should quantify L-indel–derived truncated Na_V_1.6 and WT Na_V_1.6 in DL/++ mice, truncated (DL) and full-length (D+) Na_V_1.6 in LL (DL/D+) mice, and truncated Na_V_1.6 in HP (DL/DL) mice to relate indel dose to functional channel output across genotypes. An unresolved translational question is whether allele-selective therapies could simultaneously suppress mutant *SCN8A* expression while preserving or augmenting WT *SCN8A* output—a strategy DL/++ mice are uniquely positioned to test, and that this *cis*-acting indel partially models through selective loss of DL transcript, but that has not yet been evaluated as a combined therapeutic approach. Together these comparisons define a narrow functional window for Na_V_1.6 in which both excess and deficit are poorly tolerated.

### 4.4 Genetic Rescue Validates *Scn8a*-Lowering as a Therapeutic Route

The LL phenotype is, in effect, a genetic mimic of an *Scn8a*-lowering therapeutic: a ∼30% reduction in *Scn8a* transcript converts a lethal GoF seizure phenotype into a rescued, long-lived one. This parallels the antisense-oligonucleotide work of Lenk et al. (2020), in which reducing *Scn8a* transcript by 25–50% delayed seizure onset and prolonged survival in both a GoF (R1872W) model and an *Scn1a*+/− Dravet model (24). Our allelic series provides an independent, genetic validation of transcript reduction as a strategy, and adds a quantitative caution that a pharmacological program might not otherwise reveal: the same series that shows rescue at ∼30% reduction also shows that too little reduction leaves the GoF phenotype intact (SL), whereas too much overshoots into the LoF, null-like phenotype (HP). The therapeutic target is therefore not simply “lower *Scn8a*” but “lower *Scn8a* into a defined window,” and our genotypes bracket that window experimentally.

### 4.5 Opposite Therapeutic Logic for Gain- and Loss-of-Function

A corollary of the dose model is that the two extremes of the series require opposite interventions. The SL (GoF) pole calls for suppression of Na_V_1.6 activity or expression, consistent with the clinical observation that sodium-channel blockers are preferentially effective in GoF *SCN8A* epilepsy (18), whereas the HP (LoF) pole would instead require restoration of Na_V_1.6. That a single gene can demand diametrically opposed therapeutic logic depending on dosage underscores why functional classification of variants is a prerequisite to treatment, and why a modifier that moves a patient along this axis is of direct translational interest. The molecular programs that distinguish these poles are dissected at the pathway level in the companion paper (Hammer and Bahramnejad, in prep).

### 4.6 A Graded Molecular Axis from Genotype to Transcriptional Program

By the mammalian NMD rule, PTC positioned more than ∼50–55 nucleotides upstream of the 3′-most exon–exon junction are recognized as NMD substrates (15, 16, 35). The indel variant in our colony introduces a PTC ∼26 kb upstream of that junction and is therefore predicted to engage this pathway. Read-level analysis at the frameshift indel showed selective reduction of L-bearing transcripts with preserved non-L read support and intact genomic dosage.

A related alternative is nonsense-associated altered splicing (NAS), in which a PTC can disrupt exonic splicing regulatory elements or otherwise alter splice-site choice on the mutant allele, promoting skipping of the PTC-bearing exon and reducing reads at the indel site (36, 37). Unlike NMD, NAS does not degrade mRNA but instead shifts production toward isoforms that omit the offending exon; such skipped transcripts can still contribute to total gene-level counts. Because we observe graded reduction of total *Scn8a* mRNA with increasing “L” copy number, NAS alone is an unlikely primary explanation for our data. A further PTC-linked possibility is nonsense-mediated transcriptional gene silencing (NMTGS), in which PTC recognition has been proposed to repress transcription through local chromatin remodeling. However, NMTGS has been documented mainly for immunoglobulin minigenes and was not observed for several other NMD substrates (36, 38).

A *cis* frameshift indel that truncates Na_V_1.6 upstream of the N1768D residue, provides a parsimonious molecular axis linking genotype to Na_V_1.6 dose and, in turn, to the graded phenotype. It also supplies a framework along which downstream transcriptional responses can be ordered. We find phenotype-specific divergence in differential expression across SL, LL, and HP, and the companion paper uses this same axis to resolve the beneficial and detrimental pathway programs that accompany each genotype (Hammer and Bahramnejad, in prep). Excitatory forebrain neurons are a likely locus for these effects, given their prominent role in *SCN8A* encephalopathy (39).

### 4.7 Limitations and Conclusions

Several limitations temper these conclusions. The cross-genotype comparison carries an unavoidable dose-by-age confound, because the genotypes that reduce *Scn8a* most also shorten lifespan, so molecular differences partly reflect developmental stage as well as Na_V_1.6 dose. Seizure metrics were available for a subset of mice, the rescued and heterozygous cohorts are modest in size, and a small number of source records required de-duplication. Finally, the precise quantitative mapping between transcript reduction and phenotype is likely to be species- and context-sensitive, so the ∼30% “rescue” level should be read as illustrative of a window rather than a transferable threshold. With these caveats, our data establish that a de novo, *cis*-acting LoF variant within *Scn8a* acts as a dose-dependent modifier of the N1768D GoF allele, and that the resulting allelic series both validates *Scn8a*-lowering as a therapeutic strategy and defines the functional window within which such a strategy must operate.

## Data Availability

The data that support the findings of the present study are available from the corresponding author on reasonable request. RNA-seq data supporting this study have been deposited in Figshare and are available at https://doi.org/10.6084/m9.figshare.33023150.

## Competing Interests

The authors declare that there are no competing interests associated with the manuscript.

## Funding

This work was supported by the Shay Emma Hammer Research Foundation. Hippocampal RNA-seq was supported by an award from the University of Arizona ORP Core Facilities Pilot Program.

## CRediT Author Contribution

**Erfan Bahramnejad:** Conceptualization, Data curation, Formal analysis, Methodology, Visualization, Investigation, Project administration, Writing—original draft. **Michael F. Hammer:** Conceptualization, Resources, Writing—review & editing, Funding acquisition, Methodology, Validation. **Victor J. Hiller:** Data curation, Formal analysis, Project administration.

## Ethics statement

Animal work was approved by the University of Arizona IACUC (protocol #16-160), performed in ARRIVE-compliant accredited facilities, and mice were euthanized at endpoint by cervical dislocation.

## Acknowledgment

This study was primarily supported by the Shay Emma Hammer Research Foundation. RNA-seq was funded by a Core Facility Pilot Program award from the Office of Research and Partnerships at the University of Arizona. We thank the Arizona Genetics Core and the Arizona Genomics Institute for assistance with RNA-seq and whole-genome sequencing, respectively. We also thank Taylor Camacho for assistance with animal toe clipping and genotyping.

## Supplementary Material

**Table S1.** Lifespan of male SL and LL mice with D/+ and D/D genotypes on a 98.4% C3H.B6 background.

| Mouse ID | Sex | Genotype | Age at death or Euthanasia | Date of birth | Date of birth | Date of death or euthanasia | Group |
| --- | --- | --- | --- | --- | --- | --- | --- |
| 1674 | M | HOM | 52 | 7/17/2018 | 9/7/2018 | 98.4 | LL |
| 1760 | M | HOM | 45 | 9/21/2018 | 11/5/2018 | 98.4 | LL |
| 1856 | M | HOM | 40 | 11/8/2018 | 12/18/2018 | 98.4 | LL |
| 1891 | M | HOM | 40 | 11/8/2018 | 12/18/2018 | 98.4 | LL |
| 1894 | M | HOM | 40 | 11/8/2018 | 12/18/2018 | 98.4 | LL |
| 1942 | M | HOM | 42 | 12/17/2018 | 1/28/2019 | 98.4 | LL |
| 2018 | M | HOM | 57 | 3/25/2019 | 5/21/2019 | 98.4 | LL |
| 2020 | M | HOM | 57 | 3/25/2019 | 5/21/2019 | 98.4 | LL |
| 2052 | M | HOM | 52 | 3/26/2019 | 5/17/2019 | 98.4 | LL |
| 2054 | M | HOM | 56 | 3/26/2019 | 5/21/2019 | 98.4 | LL |
| 2084 | M | HOM | 40 | 5/17/2019 | 6/26/2019 | 98.4 | LL |
| 2116 | M | HOM | 22 | 7/2/2019 | 7/24/2019 | 98.4 | SL |
| 2204 | M | HOM | 20 | 9/26/2019 | 10/16/2019 | 98.4 | SL |
| 2319 | M | HOM | 17 | 10/29/2019 | 11/15/2019 | 98.4 | SL |
| 2412 | M | HOM | 20 | 12/11/2019 | 12/31/2019 | 98.4 | SL |
| 2532 | M | HOM | 21 | 1/14/2020 | 2/4/2020 | 98.4 | SL |
| 2533 | M | HOM | 21 | 1/14/2020 | 2/4/2020 | 98.4 | SL |
| 2653 | M | HOM | 21 | 4/3/2020 | 4/24/2020 | 98.4 | SL |
| 2660 | M | HOM | 20 | 4/4/2020 | 4/24/2020 | 98.4 | SL |
| 2699 | M | HOM | 43 | 4/12/2020 | 5/25/2020 | 98.4 | LL |
| 2700 | M | HOM | 43 | 4/12/2020 | 5/25/2020 | 98.4 | LL |
| 2750 | M | HOM | 32 | 6/13/2020 | 7/15/2020 | 98.4 | LL |
| 2771 | M | HOM | 20 | 6/23/2020 | 7/13/2020 | 98.4 | SL |
| 2773 | M | HOM | 20 | 6/23/2020 | 7/13/2020 | 98.4 | SL |
| 2779 | M | HOM | 25 | 6/27/2020 | 7/22/2020 | 98.4 | SL |
| 2805 | M | HOM | 74 | 7/4/2020 | 9/16/2020 | 98.4 | LL |
| 2806 | M | HOM | 74 | 7/4/2020 | 9/16/2020 | 98.4 | LL |
| 2808 | M | HOM | 80 | 7/4/2020 | 9/22/2020 | 98.4 | LL |
| 2812 | M | HOM | 18 | 7/6/2020 | 7/24/2020 | 98.4 | SL |
| 2817 | M | HOM | 18 | 7/6/2020 | 7/24/2020 | 98.4 | SL |
| 2846 | M | HOM | 19 | 7/19/2020 | 8/7/2020 | 98.4 | SL |
| 2888 | M | HOM | 20 | 8/5/2020 | 8/25/2020 | 98.4 | SL |
| 2982 | M | HOM | 97 | 9/22/2020 | 12/28/2020 | 98.4 | LL |
| 2998 | M | HOM | 23 | 10/8/2020 | 10/31/2020 | 98.4 | SL |
| 3006 | M | HOM | 153 | 10/8/2020 | 3/10/2021 | 98.4 | LL |
| 3025 | M | HOM | 108 | 10/15/2020 | 1/31/2021 | 98.4 | LL |
| 3031 | M | HOM | 66 | 10/15/2020 | 12/20/2020 | 98.4 | LL |
| 3032 | M | HOM | 106 | 10/15/2020 | 1/29/2021 | 98.4 | LL |
| 3035 | M | HOM | 75 | 10/15/2020 | 12/29/2020 | 98.4 | LL |
| 3068 | M | HOM | 21 | 11/2/2020 | 11/23/2020 | 98.4 | SL |
| 3113 | M | HOM | 70 | 11/20/2020 | 1/29/2021 | 98.4 | LL |
| 3123 | M | HOM | 21 | 11/25/2020 | 12/16/2020 | 98.4 | SL |
| 3157 | M | HOM | 89 | 12/13/2020 | 3/12/2021 | 98.4 | LL |
| 3171 | M | HOM | 74 | 12/20/2020 | 3/4/2021 | 98.4 | LL |
| 3219 | M | HOM | 22 | 12/31/2020 | 1/22/2021 | 98.4 | SL |
| 3226 | M | HOM | 21 | 1/3/2021 | 1/24/2021 | 98.4 | SL |
| 3249 | M | HOM | 20 | 1/10/2021 | 1/30/2021 | 98.4 | SL |
| 3262 | M | HOM | 72 | 1/15/2021 | 3/28/2021 | 98.4 | LL |
| 3270 | M | HOM | 18 | 1/15/2021 | 2/2/2021 | 98.4 | SL |
| 3295 | M | HOM | 58 | 1/24/2021 | 3/23/2021 | 98.4 | LL |
| 3334 | M | HOM | 19 | 2/4/2021 | 2/23/2021 | 98.4 | SL |
| 3335 | M | HOM | 21 | 2/4/2021 | 2/25/2021 | 98.4 | SL |
| 3336 | M | HOM | 19 | 2/4/2021 | 2/23/2021 | 98.4 | SL |
| 3341 | M | HOM | 51 | 2/8/2021 | 3/31/2021 | 98.4 | LL |
| 3342 | M | HOM | 19 | 2/8/2021 | 2/27/2021 | 98.4 | SL |
| 3343 | M | HOM | 48 | 2/8/2021 | 3/28/2021 | 98.4 | LL |
| 3386 | M | HOM | 92 | 2/20/2021 | 5/23/2021 | 98.4 | LL |
| 3387 | M | HOM | 46 | 2/20/2021 | 4/7/2021 | 98.4 | LL |
| 3396 | M | HOM | 95 | 2/21/2021 | 5/27/2021 | 98.4 | LL |
| 3397 | M | HOM | 132 | 2/21/2021 | 7/3/2021 | 98.4 | LL |
| 3406 | M | HOM | 37 | 2/22/2021 | 3/31/2021 | 98.4 | LL |
| 3408 | M | HOM | 37 | 2/22/2021 | 3/31/2021 | 98.4 | LL |
| 3415 | M | HOM | 37 | 2/22/2021 | 3/31/2021 | 98.4 | LL |
| 3432 | M | HOM | 89 | 3/2/2021 | 5/30/2021 | 98.4 | LL |
| 3452 | M | HOM | 63 | 3/14/2021 | 5/16/2021 | 98.4 | LL |
| 3453 | M | HOM | 57 | 3/14/2021 | 5/10/2021 | 98.4 | LL |
| 3496 | M | HOM | 62 | 4/11/2021 | 6/12/2021 | 98.4 | LL |
| 3531 | M | HOM | 80 | 4/20/2021 | 7/9/2021 | 98.4 | LL |
| 3532 | M | HOM | 59 | 4/20/2021 | 6/18/2021 | 98.4 | LL |
| 3567 | M | HOM | 84 | 5/6/2021 | 7/29/2021 | 98.4 | LL |
| 3568 | M | HOM | 22 | 5/6/2021 | 5/28/2021 | 98.4 | SL |
| 3580 | M | HOM | 24 | 5/18/2021 | 6/11/2021 | 98.4 | SL |
| 3608 | M | HOM | 60 | 6/10/2021 | 8/9/2021 | 98.4 | LL |
| 3612 | M | HOM | 62 | 6/10/2021 | 8/11/2021 | 98.4 | LL |
| 3616 | M | HOM | 65 | 6/10/2021 | 8/14/2021 | 98.4 | LL |
| 3631 | M | HOM | 66 | 6/24/2021 | 8/29/2021 | 98.4 | LL |
| 3640 | M | HOM | 22 | 6/24/2021 | 7/16/2021 | 98.4 | SL |
| 3712 | M | HOM | 69 | 8/11/2021 | 10/19/2021 | 98.4 | LL |
| 3742 | M | HOM | 79 | 8/25/2021 | 11/12/2021 | 98.4 | LL |
| 3785 | M | HOM | 72 | 9/15/2021 | 11/26/2021 | 98.4 | LL |
| 3823 | M | HOM | 77 | 9/22/2021 | 12/8/2021 | 98.4 | LL |
| 3830 | M | HOM | 59 | 9/27/2021 | 11/25/2021 | 98.4 | LL |
| 3861 | M | HOM | 100 | 10/15/2021 | 1/23/2022 | 98.4 | LL |
| 3866 | M | HOM | 95 | 10/15/2021 | 1/18/2022 | 98.4 | LL |
| 3868 | M | HOM | 81 | 10/15/2021 | 1/4/2022 | 98.4 | LL |
| 3890 | M | HOM | 83 | 10/29/2021 | 1/20/2022 | 98.4 | LL |
| 3907 | M | HOM | 66 | 11/4/2021 | 1/9/2022 | 98.4 | LL |
| 3940 | M | HOM | 77 | 11/26/2021 | 2/11/2022 | 98.4 | LL |
| 3955 | M | HOM | 59 | 11/30/2021 | 1/28/2022 | 98.4 | LL |
| 4017 | M | HOM | 76 | 1/8/2022 | 3/25/2022 | 98.4 | LL |
| 4086 | M | HOM | 89 | 2/18/2022 | 5/18/2022 | 98.4 | LL |
| 4087 | M | HOM | 68 | 2/18/2022 | 4/27/2022 | 98.4 | LL |
| 4090 | M | HOM | 68 | 2/18/2022 | 4/27/2022 | 98.4 | LL |
| 4120 | M | HOM | 86 | 2/27/2022 | 5/24/2022 | 98.4 | LL |
| 4201 | M | HOM | 59 | 4/6/2022 | 6/4/2022 | 98.4 | LL |
| 4218 | M | HOM | 136 | 4/27/2022 | 9/10/2022 | 98.4 | LL |
| 4260 | M | HOM | 34 | 5/21/2022 | 6/24/2022 | 98.4 | LL |
| 4314 | M | HOM | 114 | 6/24/2022 | 10/16/2022 | 98.4 | LL |
| 4321 | M | HOM | 20 | 7/2/2022 | 7/22/2022 | 98.4 | SL |
| 4354 | M | HOM | 78 | 7/29/2022 | 10/15/2022 | 98.4 | LL |
| 4356 | M | HOM | 68 | 7/29/2022 | 10/5/2022 | 98.4 | LL |
| 4421 | M | HOM | 24 | 8/30/2022 | 9/23/2022 | 98.4 | SL |
| 4452 | M | HOM | 21 | 9/28/2022 | 10/19/2022 | 98.4 | SL |
| 4550 | M | HOM | 61 | 12/1/2022 | 1/31/2023 | 98.4 | LL |
| 4571 | M | HOM | 20 | 12/5/2022 | 12/25/2022 | 98.4 | SL |
| 4631 | M | HOM | 22 | 1/11/2023 | 2/2/2023 | 98.4 | SL |
| 4669 | M | HOM | 20 | 1/22/2023 | 2/11/2023 | 98.4 | SL |
| 4672 | M | HOM | 20 | 1/22/2023 | 2/11/2023 | 98.4 | SL |
| 4673 | M | HOM | 20 | 1/22/2023 | 2/11/2023 | 98.4 | SL |
| 4684 | M | HOM | 62 | 2/6/2023 | 4/9/2023 | 98.4 | LL |
| 4695 | M | HOM | 85 | 2/11/2023 | 5/7/2023 | 98.4 | LL |
| 4702 | M | HOM | 20 | 2/11/2023 | 3/3/2023 | 98.4 | SL |
| 4707 | M | HOM | 23 | 3/3/2023 | 3/26/2023 | 98.4 | SL |
| 4717 | M | HOM | 23 | 3/3/2023 | 3/26/2023 | 98.4 | SL |
| 4722 | M | HOM | 21 | 3/5/2023 | 3/26/2023 | 98.4 | SL |
| 4736 | M | HOM | 21 | 3/5/2023 | 3/26/2023 | 98.4 | SL |
| 4740 | M | HOM | 21 | 3/5/2023 | 3/26/2023 | 98.4 | SL |
| 4774 | M | HOM | 26 | 3/28/2023 | 4/23/2023 | 98.4 | SL |
| 4783 | M | HOM | 27 | 4/3/2023 | 4/30/2023 | 98.4 | LL |
| 4800 | M | HOM | 21 | 4/17/2023 | 5/8/2023 | 98.4 | SL |
| 4809 | M | HOM | 96 | 4/18/2023 | 7/23/2023 | 98.4 | LL |
| 4818 | M | HOM | 19 | 4/18/2023 | 5/7/2023 | 98.4 | SL |
| 4819 | M | HOM | 19 | 4/18/2023 | 5/7/2023 | 98.4 | SL |
| 4825 | M | HOM | 93 | 4/28/2023 | 7/30/2023 | 98.4 | LL |
| 4838 | M | HOM | 52 | 5/29/2023 | 7/20/2023 | 98.4 | LL |
| 4839 | M | HOM | 76 | 5/29/2023 | 8/13/2023 | 98.4 | LL |
| 4858 | M | HOM | 19 | 5/30/2023 | 6/18/2023 | 98.4 | SL |
| 4877 | M | HOM | 67 | 6/21/2023 | 8/27/2023 | 98.4 | LL |
| 4888 | M | HOM | 68 | 6/30/2023 | 9/6/2023 | 98.4 | LL |
| 4893 | M | HOM | 76 | 6/30/2023 | 9/14/2023 | 98.4 | LL |
| 4920 | M | HOM | 51 | 7/26/2023 | 9/15/2023 | 98.4 | LL |
| 4922 | M | HOM | 67 | 7/26/2023 | 10/1/2023 | 98.4 | LL |
| 4965 | M | HOM | 61 | 8/23/2023 | 10/23/2023 | 98.4 | LL |
| 4966 | M | HOM | 53 | 8/23/2023 | 10/15/2023 | 98.4 | LL |
| 4973 | M | HOM | 63 | 8/22/2023 | 10/24/2023 | 98.4 | LL |
| 4984 | M | HOM | 21 | 8/28/2023 | 9/18/2023 | 98.4 | SL |
| 4985 | M | HOM | 21 | 8/28/2023 | 9/18/2023 | 98.4 | SL |
| 4992 | M | HOM | 78 | 9/10/2023 | 11/27/2023 | 98.4 | LL |
| 5000 | M | HOM | 91 | 9/15/2023 | 12/15/2023 | 98.4 | LL |
| 5084 | M | HOM | 72 | 10/23/2023 | 1/3/2024 | 98.4 | LL |
| 5127 | M | HOM | 57 | 11/15/2023 | 1/11/2024 | 98.4 | LL |
| 5134 | M | HOM | 62 | 11/17/2023 | 1/18/2024 | 98.4 | LL |
| 5136 | M | HOM | 55 | 11/17/2023 | 1/11/2024 | 98.4 | LL |
| 5144 | M | HOM | 51 | 11/19/2023 | 1/9/2024 | 98.4 | LL |
| 5146 | M | HOM | 62 | 11/19/2023 | 1/20/2024 | 98.4 | LL |
| 5154 | M | HOM | 20 | 11/20/2023 | 12/10/2023 | 98.4 | SL |
| 5177 | M | HOM | 80 | 12/8/2023 | 2/26/2024 | 98.4 | LL |
| 5184 | M | HOM | 78 | 12/10/2023 | 2/26/2024 | 98.4 | LL |
| 5216 | M | HOM | 34 | 12/30/2023 | 2/2/2024 | 98.4 | LL |
| 5261 | M | HOM | 25 | 1/16/2024 | 2/10/2024 | 98.4 | SL |
| 5290 | M | HOM | 47 | 1/25/2024 | 3/12/2024 | 98.4 | LL |
| 5315 | M | HOM | 20 | 2/17/2024 | 3/8/2024 | 98.4 | SL |
| 5354 | M | HOM | 52 | 3/1/2024 | 4/22/2024 | 98.4 | LL |
| 5380 | M | HOM | 54 | 3/7/2024 | 4/30/2024 | 98.4 | LL |
| 5395 | M | HOM | 61 | 3/19/2024 | 5/19/2024 | 98.4 | LL |
| 5398 | M | HOM | 56 | 3/19/2024 | 5/14/2024 | 98.4 | LL |
| 5416 | M | HOM | 54 | 4/24/2024 | 6/17/2024 | 98.4 | LL |
| 5417 | M | HOM | 53 | 4/24/2024 | 6/16/2024 | 98.4 | LL |
| 5423 | M | HOM | 74 | 4/20/2024 | 7/3/2024 | 98.4 | LL |
| 5435 | M | HOM | 39 | 4/28/2024 | 6/6/2024 | 98.4 | LL |
| 5437 | M | HOM | 39 | 4/28/2024 | 6/6/2024 | 98.4 | LL |
| 5438 | M | HOM | 65 | 4/28/2024 | 7/2/2024 | 98.4 | LL |
| 5450 | M | HOM | 33 | 5/4/2024 | 6/6/2024 | 98.4 | LL |
| 5493 | M | HOM | 38 | 6/4/2024 | 7/12/2024 | 98.4 | LL |
| 5498 | M | HOM | 26 | 6/7/2024 | 7/3/2024 | 98.4 | LL |
| 5499 | M | HOM | 26 | 6/7/2024 | 7/3/2024 | 98.4 | LL |
| 5502 | M | HOM | 35 | 6/7/2024 | 7/12/2024 | 98.4 | LL |
| 5508 | M | HOM | 26 | 6/7/2024 | 7/3/2024 | 98.4 | LL |
| 5510 | M | HOM | 32 | 6/7/2024 | 7/9/2024 | 98.4 | LL |
| 5557 | M | HOM | 68 | 7/1/2024 | 9/7/2024 | 98.4 | LL |
| 5581 | M | HOM | 67 | 7/12/2024 | 9/17/2024 | 98.4 | LL |
| 5592 | M | HOM | 22 | 7/16/2024 | 8/7/2024 | 98.4 | SL |
| 5593 | M | HOM | 22 | 7/16/2024 | 8/7/2024 | 98.4 | SL |
| 5629 | M | HOM | 30 | 8/24/2024 | 9/23/2024 | 98.4 | LL |
| 5643 | M | HOM | 24 | 8/30/2024 | 9/23/2024 | 98.4 | SL |
| 5722 | M | HOM | 55 | 9/25/2024 | 11/19/2024 | 98.4 | LL |
| 5759 | M | HOM | 20 | 10/30/2024 | 11/19/2024 | 98.4 | SL |
| 5778 | M | HOM | 20 | 11/6/2024 | 11/26/2024 | 98.4 | SL |
| 5797 | M | HOM | 60 | 11/16/2024 | 1/15/2025 | 98.4 | LL |
| 5798 | M | HOM | 62 | 11/16/2024 | 1/17/2025 | 98.4 | LL |
| 5834 | M | HOM | 77 | 12/5/2024 | 2/20/2025 | 98.4 | LL |
| 5843 | M | HOM | 51 | 12/31/2024 | 2/20/2025 | 98.4 | LL |
| 5963 | M | HOM | 20 | 2/5/2025 | 2/25/2025 | 98.4 | SL |
| 6012 | M | HOM | 58 | 3/16/2025 | 5/13/2025 | 98.4 | LL |
| 6092 | M | HOM | 21 | 4/9/2025 | 4/30/2025 | 98.4 | SL |
| 6097 | M | HOM | 22 | 4/9/2025 | 5/1/2025 | 98.4 | SL |
| 6758 | M | HOM | 23 | 5/21/2026 | 6/13/2026 | 98.4 | SL |
| 6760 | M | HOM | 21 | 5/21/2026 | 6/11/2026 | 98.4 | SL |
| 2864 | M | HOM | 21 | 7/24/2020 | 8/14/2020 | 98.4 | HP |
| 2867 | M | HOM | 21 | 7/24/2020 | 8/14/2020 | 98.4 | HP |
| 2921 | M | HOM | 16 | 8/9/2020 | 8/25/2020 | 98.4 | HP |
| 3134 | M | HOM | 19 | 11/30/2020 | 12/19/2020 | 98.4 | HP |
| 3304 | M | HOM | 15 | 1/29/2021 | 2/13/2021 | 98.4 | HP |
| 3305 | M | HOM | 15 | 1/29/2021 | 2/13/2021 | 98.4 | HP |
| 3306 | M | HOM | 15 | 1/29/2021 | 2/13/2021 | 98.4 | HP |
| 3309 | M | HOM | 15 | 1/29/2021 | 2/13/2021 | 98.4 | HP |
| 3325 | M | HOM | 16 | 2/3/2021 | 2/19/2021 | 98.4 | HP |
| 3326 | M | HOM | 16 | 2/3/2021 | 2/19/2021 | 98.4 | HP |
| 3328 | M | HOM | 16 | 2/3/2021 | 2/19/2021 | 98.4 | HP |
| 3367 | M | HOM | 19 | 2/20/2021 | 3/11/2021 | 98.4 | HP |
| 3428 | M | HOM | 15 | 2/25/2021 | 3/12/2021 | 98.4 | HP |
| 3429 | M | HOM | 15 | 2/25/2021 | 3/12/2021 | 98.4 | HP |
| 3442 | M | HOM | 20 | 3/9/2021 | 3/29/2021 | 98.4 | HP |
| 3466 | M | HOM | 16 | 4/3/2021 | 4/19/2021 | 98.4 | HP |
| 3467 | M | HOM | 16 | 4/3/2021 | 4/19/2021 | 98.4 | HP |
| 3478 | M | HOM | 16 | 4/7/2021 | 4/23/2021 | 98.4 | HP |
| 3524 | M | HOM | 18 | 4/19/2021 | 5/7/2021 | 98.4 | HP |
| 3560 | M | HOM | 17 | 5/2/2021 | 5/19/2021 | 98.4 | HP |
| 3561 | M | HOM | 17 | 5/2/2021 | 5/19/2021 | 98.4 | HP |
| 3562 | M | HOM | 17 | 5/2/2021 | 5/19/2021 | 98.4 | HP |
| 3576 | M | HOM | 13 | 5/15/2021 | 5/28/2021 | 98.4 | HP |
| 3577 | M | HOM | 13 | 5/15/2021 | 5/28/2021 | 98.4 | HP |
| 3581 | M | HOM | 18 | 5/18/2021 | 6/5/2021 | 98.4 | HP |
| 3707 | M | HOM | 19 | 8/9/2021 | 8/28/2021 | 98.4 | HP |
| 3948 | M | HOM | 19 | 11/28/2021 | 12/17/2021 | 98.4 | HP |
| 3949 | M | HOM | 19 | 11/28/2021 | 12/17/2021 | 98.4 | HP |
| 3964 | M | HOM | 16 | 12/4/2021 | 12/20/2021 | 98.4 | HP |
| 4099 | M | HOM | 15 | 2/19/2022 | 3/6/2022 | 98.4 | HP |
| 4104 | M | HOM | 15 | 2/19/2022 | 3/6/2022 | 98.4 | HP |
| 4145 | M | HOM | 14 | 3/11/2022 | 3/25/2022 | 98.4 | HP |
| 4158 | M | HOM | 18 | 3/14/2022 | 4/1/2022 | 98.4 | HP |
| 4159 | M | HOM | 18 | 3/14/2022 | 4/1/2022 | 98.4 | HP |
| 4266 | M | HOM | 14 | 5/22/2022 | 6/5/2022 | 98.4 | HP |
| 4285 | M | HOM | 17 | 5/25/2022 | 6/11/2022 | 98.4 | HP |
| 4438 | M | HOM | 15 | 9/8/2022 | 9/23/2022 | 98.4 | HP |
| 4533 | M | HOM | 15 | 11/23/2022 | 12/8/2022 | 98.4 | HP |
| 5279 | M | HOM | 15 | 1/22/2024 | 2/6/2024 | 98.4 | HP |
| 5280 | M | HOM | 15 | 1/22/2024 | 2/6/2024 | 98.4 | HP |
| 5342 | M | HOM | 20 | 2/22/2024 | 3/13/2024 | 98.4 | HP |
| 5803 | M | HOM | 21 | 12/5/2024 | 12/26/2024 | 98.4 | HP |
| 6000 | M | HOM | 13 | 2/27/2025 | 3/12/2025 | 98.4 | HP |
| 6081 | M | HOM | 18 | 4/7/2025 | 4/25/2025 | 98.4 | HP |
| 6126 | M | HOM | 13 | 5/1/2025 | 5/14/2025 | 98.4 | HP |
| 6749 | M | HOM | 17 | 5/8/2026 | 5/25/2026 | 98.4 | HP |
| 6787 | M | HOM | 20 | 5/31/2026 | 6/20/2026 | 98.4 | HP |
| 6789 | M | HOM | 20 | 5/31/2026 | 6/20/2026 | 98.4 | HP |
| 6261 | M | HET | 218 | 6/27/2025 | 1/31/2026 | 98.4 | LL |
| 6263 | M | HET | 218 | 6/27/2025 | 1/31/2026 | 98.4 | LL |
| 6264 | M | HET | 218 | 6/27/2025 | 1/31/2026 | 98.4 | LL |
| 6265 | M | HET | 218 | 6/27/2025 | 1/31/2026 | 98.4 | LL |
| 6266 | M | HET | 218 | 6/27/2025 | 1/31/2026 | 98.4 | LL |
| 6409 | M | HET | 84 | 9/16/2025 | 12/9/2025 | 98.4 | LL |
| 6410 | M | HET | 112 | 9/16/2025 | 1/6/2026 | 98.4 | LL |
| 6615 | M | HET | 80 | 1/4/2026 | 3/25/2026 | 98.4 | LL |
| 6616 | M | HET | 82 | 1/4/2026 | 3/27/2026 | 98.4 | LL |
| 1605 | M | HET | 741 | 5/8/2018 | 5/18/2020 | 98.4 | LL |
| 6159 | M | HET | 72 | 5/21/2025 | 8/1/2025 | 98.4 | SL |
| 6544 | M | HET | 112 | 11/19/2025 | 3/11/2026 | 98.4 | SL |
| 6545 | M | HET | 117 | 11/19/2025 | 3/16/2026 | 98.4 | SL |
| 6641 | M | HET | 72 | 1/31/2026 | 4/13/2026 | 98.4 | SL |
| 1601 | M | HET | 98 | 5/18/2018 | 8/24/2018 | 98.4 | SL |
| 1602 | M | HET | 142 | 5/18/2018 | 10/7/2018 | 98.4 | SL |
| 1603 | M | HET | 181 | 5/18/2018 | 11/15/2018 | 98.4 | SL |
| 1604 | M | HET | 160 | 5/18/2018 | 10/25/2018 | 98.4 | SL |
| 6668 | M | HET | 108 | 2/27/2026 | 6/15/2026 | 98.4 | SL |
| 6671 | M | HET | 108 | 2/27/2026 | 6/15/2026 | 98.4 | SL |
| 3951 | M | HET | 73 | 11/30/2021 | 2/11/2022 | 98.4 | SL |
| 4681 | M | HET | 76 | 2/6/2023 | 4/23/2023 | 98.4 | SL |
| 6330 | M | HET | 77 | 7/23/2025 | 10/8/2025 | 98.4 | SL |
| 3800 | M | HET | 81 | 9/19/2021 | 12/9/2021 | 98.4 | SL |
| 3741 | M | HET | 82 | 8/25/2021 | 11/15/2021 | 98.4 | SL |
| 6549 | M | HET | 89 | 11/19/2025 | 2/16/2026 | 98.4 | SL |
| 3414 | M | HET | 99 | 2/22/2021 | 6/1/2021 | 98.4 | SL |
| 2698 | M | HET | 101 | 4/12/2020 | 7/22/2020 | 98.4 | SL |
| 2723 | M | HET | 105 | 4/28/2020 | 8/11/2020 | 98.4 | SL |
| 3024 | M | HET | 105 | 10/15/2020 | 1/28/2021 | 98.4 | SL |
| 4645 | M | HET | 106 | 1/18/2023 | 5/4/2023 | 98.4 | SL |
| 5179 | M | HET | 110 | 12/8/2023 | 3/27/2024 | 98.4 | SL |
| 3023 | M | HET | 114 | 10/15/2020 | 2/6/2021 | 98.4 | SL |
| 6640 | M | HET | 115 | 1/31/2026 | 5/26/2026 | 98.4 | SL |
| 2610 | M | HET | 119 | 2/22/2020 | 6/20/2020 | 98.4 | SL |
| 2611 | M | HET | 123 | 2/22/2020 | 6/24/2020 | 98.4 | SL |
| 3422 | M | HET | 124 | 2/22/2021 | 6/26/2021 | 98.4 | SL |
| 2946 | M | HET | 137 | 9/2/2020 | 1/17/2021 | 98.4 | SL |
| 2383 | M | HET | 150 | 11/18/2019 | 4/16/2020 | 98.4 | SL |

**Table S2.** Age of seizure onset and seizure frequency in male SL and LL mice with D/+ and D/D genotypes on a 98.4% C3H.B6 background.

| Mouse ID | Age of onset | Total TCS | TCS/day | Sex | Group | Genotype | Background |
| --- | --- | --- | --- | --- | --- | --- | --- |
| 3031 | 64 | 86 | 28.7 | M | LL | HOM | 98.4 |
| 3032 | 54 | 115 | 2.2 | M | LL | HOM | 98.4 |
| 3035 | 54 | 108 | 5.4 | M | LL | HOM | 98.4 |
| 3113 | 65 | 99 | 5.0 | M | LL | HOM | 98.4 |
| 3171 | 44 | 73 | 2.4 | M | LL | HOM | 98.4 |
| 3432 | 42 | 67 | 1.5 | M | LL | HOM | 98.4 |
| 3453 | 41 | 16 | 0.7 | M | LL | HOM | 98.4 |
| 3616 | 41 | 59 | 2.5 | M | LL | HOM | 98.4 |
| 3785 | 43 | 100 | 3.4 | M | LL | HOM | 98.4 |
| 3830 | 38 | 85 | 4.3 | M | LL | HOM | 98.4 |
| 6238 | 19 | 22 | 11.0 | M | SL | HOM | 98.4 |
| 6240 | 20 | 34 | 34.0 | M | SL | HOM | 98.4 |
| 6241 | 19 | 17 | 8.5 | M | SL | HOM | 98.4 |
| 6314 | 17 | 22 | 5.5 | M | SL | HOM | 98.4 |
| 6316 | 18 | 17 | 8.5 | M | SL | HOM | 98.4 |
| 6261 | NA | 0 | 0 | M | LL | HET | 98.4 |
| 6263 | NA | 0 | 0 | M | LL | HET | 98.4 |
| 6264 | NA | 0 | 0 | M | LL | HET | 98.4 |
| 6265 | NA | 0 | 0 | M | LL | HET | 98.4 |
| 6266 | NA | 0 | 0 | M | LL | HET | 98.4 |

**Table S3. Candidate variants from WhatsHap-phased trio WGS passing genotype filter rules near *Scn8a*-N1768D**

Full table is provided as Excel file Table S3.xlsx.

**Table S4.** RNA-seq REF and ALT read counts at the *Scn8a* frameshift indel in WT, SL, LL, and HP mice.

| Sample id | group | pos | ref | alt | Ref reads | Alt reads | Ambiguous reads | Other pattern reads | Skipped reads | Informative Ref or alt only | alt_fraction informative | Expected genotype | Called genotype | Dosage score |
| --- | --- | --- | --- | --- | --- | --- | --- | --- | --- | --- | --- | --- | --- | --- |
| 6138 | SL | 100911188 | CGT | C | 148 | 0 | 0 | 11 | 0 | 148 | 0.0 | ref/ref | ref/ref | 0 |
| 6139 | SL | 100911188 | CGT | C | 257 | 0 | 0 | 25 | 0 | 257 | 0.0 | ref/ref | ref/ref | 0 |
| 6153 | SL | 100911188 | CGT | C | 217 | 0 | 0 | 21 | 0 | 217 | 0.0 | ref/ref | ref/ref | 0 |
| 6170 | LL | 100911188 | CGT | C | 62 | 10 | 0 | 11 | 0 | 72 | 0.138889 | ref/del | ref/del | 1 |
| 6171 | LL | 100911188 | CGT | C | 115 | 18 | 0 | 20 | 0 | 133 | 0.135338 | ref/del | ref/del | 1 |
| 6209 | LL | 100911188 | CGT | C | 118 | 17 | 0 | 13 | 0 | 135 | 0.125926 | ref/del | ref/del | 1 |
| 6334 | HP | 100911188 | CGT | C | 0 | 40 | 0 | 15 | 1 | 40 | 1.0 | del/del | del/del | 2 |
| 6339 | HP | 100911188 | CGT | C | 0 | 35 | 0 | 8 | 0 | 35 | 1.0 | del/del | del/del | 2 |
| 6431 | HP | 100911188 | CGT | C | 0 | 28 | 0 | 15 | 0 | 28 | 1.0 | del/del | del/del | 2 |
| 6336 | WT | 100911188 | CGT | C | 191 | 0 | 0 | 13 | 5 | 191 | 0.0 | ref/ref | ref/ref | 0 |
| 6338 | WT | 100911188 | CGT | C | 244 | 0 | 0 | 10 | 1 | 244 | 0.0 | ref/ref | ref/ref | 0 |
| 6465 | WT | 100911188 | CGT | C | 194 | 0 | 0 | 24 | 2 | 194 | 0.0 | ref/ref | ref/ref | 0 |

**Video S1.** Hindlimb paralysis phenotype in homozygous HP (DL/DL) *Scn8a*-N1768D mice on the 98.4% C3H·B6 background.

**Video S2.** Audiogenic seizure test in heterozygous D/+ SL and D/+ LL mice.

**Video S3.** Genomic context of the *Scn8a* 2-bp coding frameshift relative to N1768D and predicted truncation of Na_V_1.6 at codon 914.

## Notes

https://doi.org/10.6084/m9.figshare.33023150

### Competing Interest Statement

The authors have declared no competing interest.

## References

1. Catterall WA. Forty Years of Sodium Channels: Structure, Function, Pharmacology, and Epilepsy. Neurochem Res. 2017;42(9):2495–504.

2. Hammer MF. Transgenic mouse models of sodium and potassium channelopathies in epilepsy: insights into disease mechanisms and therapeutics. Biosci Rep. 2025;45(10).

3. Hammer MF, Xia M, Schreiber JM. SCN8A-related epilepsy and/or neurodevelopmental disorders. GeneReviews®[Internet]. 2023.

4. Villapol S, Janatpour ZC, Affram KO, Symes AJ. The Renin Angiotensin System as a Therapeutic Target in Traumatic Brain Injury. Neurotherapeutics. 2023;20(6):1565–91.

5. Ayoub D, Jaafar F, Al-Hajje A, Salameh P, Jost J, Hmaimess G, et al. Predictors of drug-resistant epilepsy in childhood epilepsy syndromes: A subgroup analysis from a prospective cohort study. Epilepsia. 2024;65(10):2995–3009.

6. Consortium SAR. A research roadmap for SCN8A-related disorders: addressing knowledge gaps and aligning research priorities across stakeholders. Orphanet J Rare Dis. 2025;20(1):444.

7. Genin E, Feingold J, Clerget-Darpoux F. Identifying modifier genes of monogenic disease: strategies and difficulties. Human genetics. 2008;124:357–68.

8. Rahit K, Tarailo-Graovac M. Genetic Modifiers and Rare Mendelian Disease. Genes (Basel). 2020;11(3).

9. Meisler MH, O’Brien JE. Gene Interactions and Modifiers in Epilepsy. In: Noebels JL, Avoli M, Rogawski MA, Olsen RW, Delgado-Escueta AV, editors. Jasper’s Basic Mechanisms of the Epilepsies. 4th ed. Bethesda (MD) 2012.

10. Yu W, Hill SF, Xenakis JG, Pardo-Manuel de Villena F, Wagnon JL, Meisler MH. Gabra2 is a genetic modifier of Scn8a encephalopathy in the mouse. Epilepsia. 2020;61(12):2847–56.

11. Jones JM, Meisler MH. Modeling human epilepsy by TALEN targeting of mouse sodium channel Scn8a. genesis. 2014;52(2):141–8.

12. Bahramnejad E, Barney ER, Lester S, Hurtado A, Thompson T, Watkins JC, et al. Greater female than male resilience to mortality and morbidity in the Scn8a mouse model of pediatric epilepsy. International Journal of Neuroscience. 2024;134(12):1611–23.

13. Wengert ER, Wenker IC, Wagner EL, Wagley PK, Gaykema RP, Shin JB, et al. Adrenergic Mechanisms of Audiogenic Seizure-Induced Death in a Mouse Model of SCN8A Encephalopathy. Front Neurosci. 2021;15:581048.

14. Fang H, Wu Y, Narzisi G, O’Rawe JA, Barron LT, Rosenbaum J, et al. Reducing INDEL calling errors in whole genome and exome sequencing data. Genome Med. 2014;6(10):89.

15. Schweingruber C, Rufener SC, Zund D, Yamashita A, Muhlemann O. Nonsense-mediated mRNA decay - mechanisms of substrate mRNA recognition and degradation in mammalian cells. Biochim Biophys Acta. 2013;1829(6-7):612–23.

16. Maquat LE. Nonsense-mediated mRNA decay in mammals. J Cell Sci. 2005;118(Pt 9):1773–6.

17. O’Brien JE, Meisler MH. Sodium channel SCN8A (Nav1. 6): properties and de novo mutations in epileptic encephalopathy and intellectual disability. Frontiers in genetics. 2013;4:213.

18. Johannesen KM, Liu Y, Koko M, Gjerulfsen CE, Sonnenberg L, Schubert J, et al. Genotype-phenotype correlations in SCN8A-related disorders reveal prognostic and therapeutic implications. Brain. 2022;145(9):2991–3009.

19. Meisler MH, Hill SF, Yu W. Sodium channelopathies in neurodevelopmental disorders. Nat Rev Neurosci. 2021;22(3):152–66.

20. Larsen J, Carvill GL, Gardella E, Kluger G, Schmiedel G, Barisic N, et al. The phenotypic spectrum of SCN8A encephalopathy. Neurology. 2015;84(5):480–9.

21. Hammer MF, Wagnon JL, Mefford HC, Meisler MH. SCN8A-related epilepsy with encephalopathy. 2016.

22. Chung KM, Hack J, Andrews J, Galindo-Kelly M, Schreiber J, Watkins J, et al. Clinical severity is correlated with age at seizure onset and biophysical properties of recurrent gain of function variants associated with SCN8A-related epilepsy. Epilepsia. 2023;64(12):3365–76.

23. Wagnon JL, Korn MJ, Parent R, Tarpey TA, Jones JM, Hammer MF, et al. Convulsive seizures and SUDEP in a mouse model of SCN8A epileptic encephalopathy. Hum Mol Genet. 2015;24(2):506–15.

24. Lenk GM, Jafar-Nejad P, Hill SF, Huffman LD, Smolen CE, Wagnon JL, et al. Scn8a antisense oligonucleotide is protective in mouse models of SCN8A encephalopathy and Dravet syndrome. Annals of neurology. 2020;87(3):339–46.

25. Miller AR, Hawkins NA, McCollom CE, Kearney JA. Mapping genetic modifiers of survival in a mouse model of Dravet syndrome. Genes Brain Behav. 2014;13(2):163–72.

26. Hawkins NA, Nomura T, Duarte S, Barse L, Williams RW, Homanics GE, et al. Gabra2 is a genetic modifier of Dravet syndrome in mice. Mammalian Genome. 2021;32(5):350–63.

27. Trudeau MM, Dalton JC, Day JW, Ranum LP, Meisler MH. Heterozygosity for a protein truncation mutation of sodium channel SCN8A in a patient with cerebellar atrophy, ataxia, and mental retardation. J Med Genet. 2006;43(6):527–30.

28. Johannesen KM, Gardella E, Encinas AC, Lehesjoki AE, Linnankivi T, Petersen MB, et al. The spectrum of intermediate SCN8A-related epilepsy. Epilepsia. 2019;60(5):830–44.

29. Hack JB, Horning K, Juroske Short DM, Schreiber JM, Watkins JC, Hammer MF. Distinguishing Loss-of-Function and Gain-of-Function SCN8A Variants Using a Random Forest Classification Model Trained on Clinical Features. Neurol Genet. 2023;9(3):e200060.

30. Inglis GAS, Wong JC, Butler KM, Thelin JT, Mistretta OC, Wu X, et al. Mutations in the Scn8a DIIS4 voltage sensor reveal new distinctions among hypomorphic and null Na(v) 1.6 sodium channels. Genes Brain Behav. 2020;19(4):e12612.

31. Yu W, Hill SF, Huang Y, Zhu L, Demetriou Y, Ziobro J, et al. Allele-Specific Editing of a Dominant SCN8A Epilepsy Variant Protects against Seizures and Lethality in a Murine Model. Ann Neurol. 2024;96(5):958–69.

32. Al Saneh A, Arrieta MFH, Gissot L, O’Bryan M, Li R, Buchanan G, et al. Nonsense-mediated decay influences position-dependent effects of SCN2A premature stop codons on neuronal excitability and behavior. Mol Psychiatry. 2026.

33. Wagnon JL, Barker BS, Ottolini M, Park Y, Volkheimer A, Valdez P, et al. Loss-of-function variants of SCN8A in intellectual disability without seizures. Neurology: Genetics. 2017;3(4):e170.

34. Burgess DL, Kohrman DC, Galt J, Plummer NW, Jones JM, Spear B, et al. Mutation of a new sodium channel gene, Scn8a, in the mouse mutant ‘motor endplate disease’. Nat Genet. 1995;10(4):461–5.

35. Nagy E, Maquat LE. A rule for termination-codon position within intron-containing genes: when nonsense affects RNA abundance. Trends Biochem Sci. 1998;23(6):198–9.

36. Hwang J, Kim YK. When a ribosome encounters a premature termination codon. BMB reports. 2013;46(1):9.

37. Wang J, Chang YF, Hamilton JI, Wilkinson MF. Nonsense-associated altered splicing: a frame-dependent response distinct from nonsense-mediated decay. Mol Cell. 2002;10(4):951–7.

38. Buhler M, Mohn F, Stalder L, Muhlemann O. Transcriptional silencing of nonsense codon-containing immunoglobulin minigenes. Mol Cell. 2005;18(3):307–17.

39. Bunton-Stasyshyn RKA, Wagnon JL, Wengert ER, Barker BS, Faulkner A, Wagley PK, et al. Prominent role of forebrain excitatory neurons in SCN8A encephalopathy. Brain. 2019;142(2):362–75.

